# Immunometabolic IRG1–NLRC5 Axis Controls Inflammatory Cell Death, PANoptosis

**DOI:** 10.64898/2026.11.19.765772

**Authors:** Nagakannan Pandian, Balamurugan Sundaram, Sivakumar Prasanth Kumar, Gustavo Palacios, Alexander P. Young, Roman Sarkar, Chadi A. El Farran, Thirumala-Devi Kanneganti

## Abstract

Innate immunity is the first defense against pathogens, sterile injury, and homeostatic perturbations. One key mechanism used by the innate immune system for host defense is inflammatory cell death, PANoptosis, which eliminates infected or damaged cells and releases cytokines and other danger signals that can further propagate the immune response. Given these inflammatory functions, PANoptosis must be tightly regulated to prevent excess inflammation and pathology. Cellular metabolism is a key regulator of innate immunity, yet how metabolic reprogramming regulates innate immune signaling to control cell death, inflammation, and disease remains poorly understood. Here, we identify the immune-responsive gene 1 (IRG1)– itaconate axis as a critical regulator of PANoptosis. Innate immune triggers robustly induced IRG1 expression, leading to itaconate accumulation and driving IRG1-dependent inflammatory cell death. Mechanistically, glutaminolysis fueled IRG1-dependent itaconate synthesis, and inhibition of glutamine metabolism reduced itaconate accumulation and blocked PANoptosis. Furthermore, NAD⁺ depletion, a known inducer of NLRC5-mediated PANoptosis that has been connected to hemolytic, infectious, and inflammatory conditions, acted upstream inducing IRG1 expression and promoting NLRC5-PANoptosome assembly. Together, these findings highlight IRG1 as a metabolic node linking innate immunity to PANoptosis, identifying IRG1 as a potential therapeutic target in hemolytic and inflammatory diseases.

## MAIN

Innate immunity provides the first line of defense against infectious agents, sterile injury, and homeostatic disruptions. This defense mechanism relies on pattern recognition receptors (PRRs) to detect pathogen-associated molecular patterns (PAMPs), damage-associated molecular patterns (DAMPs), cytokine signaling, and metabolic and homeostatic changes to trigger inflammatory signaling and programmed cell death pathways, including PANoptosis^1–3^. Innate immune pathways are also closely linked to cellular metabolism. For example, PRR activation induces metabolic remodeling that can form a feedback loop to engage additional PRRs and drive cytokine release and lytic cell death^4–6^. This amplification loop between innate immune signaling, cell death, and metabolic disruptions can exacerbate cellular and tissue damage, contributing to numerous disorders, including diabetes and obesity as well as cancer, neurodegeneration, and infections^5–10^. Despite these clear connections in disease, the specific molecular mechanisms through which metabolism and innate immunity are connected and control inflammatory cell death are still not fully understood, limiting the ability to therapeutically target these processes.

Here, we identified the metabolic enzyme immune-responsive gene 1 (IRG1) as a key regulator of innate immune inflammatory cell death, PANoptosis. We found that innate immune activation triggered metabolic rewiring to induce IRG1-dependent itaconate synthesis that drove NLRC5- and NLRP12-mediated PANoptosis. Deletion of *Irg1* or glutamine restriction blocked mitochondrial reactive oxygen species (ROS) generation and prevented this cell death. Collectively, our findings identify IRG1 as a key regulator linking metabolic reprogramming with the innate immune response, highlighting IRG1 as a therapeutic target in diseases associated with PANoptosis.

### IRG1 is upregulated under hemolytic conditions

Metabolic disruptions have been linked to innate immune activation across several disease states^5–10^; however, the mechanistic connections between metabolic remodeling and inflammatory cell death remain unknown. Therefore, we sought to define this connection by focusing on pathogenic conditions where metabolic disruptions are linked to cell death. One such example is hemolytic conditions, e.g., those seen during infection, hemolytic disease, and inflammatory disorders, where mitochondrial dysfunction and metabolic alterations can enhance immune responses, inflammation, and tissue damage^11–14^. Heme is a prototypical DAMP released during hemolysis, infection, or hemorrhage. Combinations of heme with PAMPs or the cytokine TNF alter metabolic processes to deplete nicotinamide adenine dinucleotide (NAD⁺), driving NLRC5-dependent inflammatory cell death, PANoptosis^15^ (Figure S1A and S1B). PANoptosis is an innate immune, inflammatory cell death pathway initiated by innate immune sensors and driven by caspases and RIPKs through PANoptosomes^2,15–17^. Among innate immune sensors, and particularly NLRs, NLRC5 is largely understudied, and the specific mechanistic connections between metabolic changes and its activation remain unknown.

To identify potential mechanistic regulators of NLRC5 and the subsequent cell death, we performed RNAseq on wild-type (WT) bone-marrow derived macrophages (BMDMs) treated with heme plus the PAMP Pam3CSK4 (Pam3) or heme plus TNF, conditions mimicking the proinflammatory environment of hemolytic diseases and infection. Given the central role of mitochondria in regulating metabolism^18,19^, we focused on differentially expressed mitochondria-associated genes. We identified *Irg1* as one of the most significantly upregulated genes in both heme plus Pam3- or heme plus TNF-treated BMDMs (Figure 1A and Figure S1C). *Irg1*, which encodes the protein IRG1 (also known as aconitate decarboxylase 1, ACOD1), is a mitochondrial enzyme that is upregulated under inflammatory conditions and catalyzes the decarboxylation of tricarboxylic acid (TCA) intermediate cis-aconitate to generate the immunometabolite itaconate^20^. The upregulation of IRG1 was further confirmed using western blotting, where we observed a time-dependent increase in IRG1 protein expression in BMDMs treated with heme plus Pam3 (Figure 1B) and heme plus TNF (Figure 1C). To determine the contribution of individual ligands to IRG1 induction, we examined IRG1 expression following treatment with each ligand alone or in combination. Heme, TNF, or Pam3 alone induced IRG1 expression; however, the combination of heme with TNF or Pam3 resulted in greater IRG1 expression compared with the individual treatments (Figure 1D, 1E, S1D, and S1E). We also measured itaconate levels and found that heme and TNF individually increased itaconate levels, and that heme plus TNF treatment led to significantly more itaconate accumulation compared to heme and TNF treatments alone (Figure 1F). These results suggest that IRG1 may have a critical role in the metabolic regulation of the innate immune response to hemolytic triggers.

**Figure 1.**
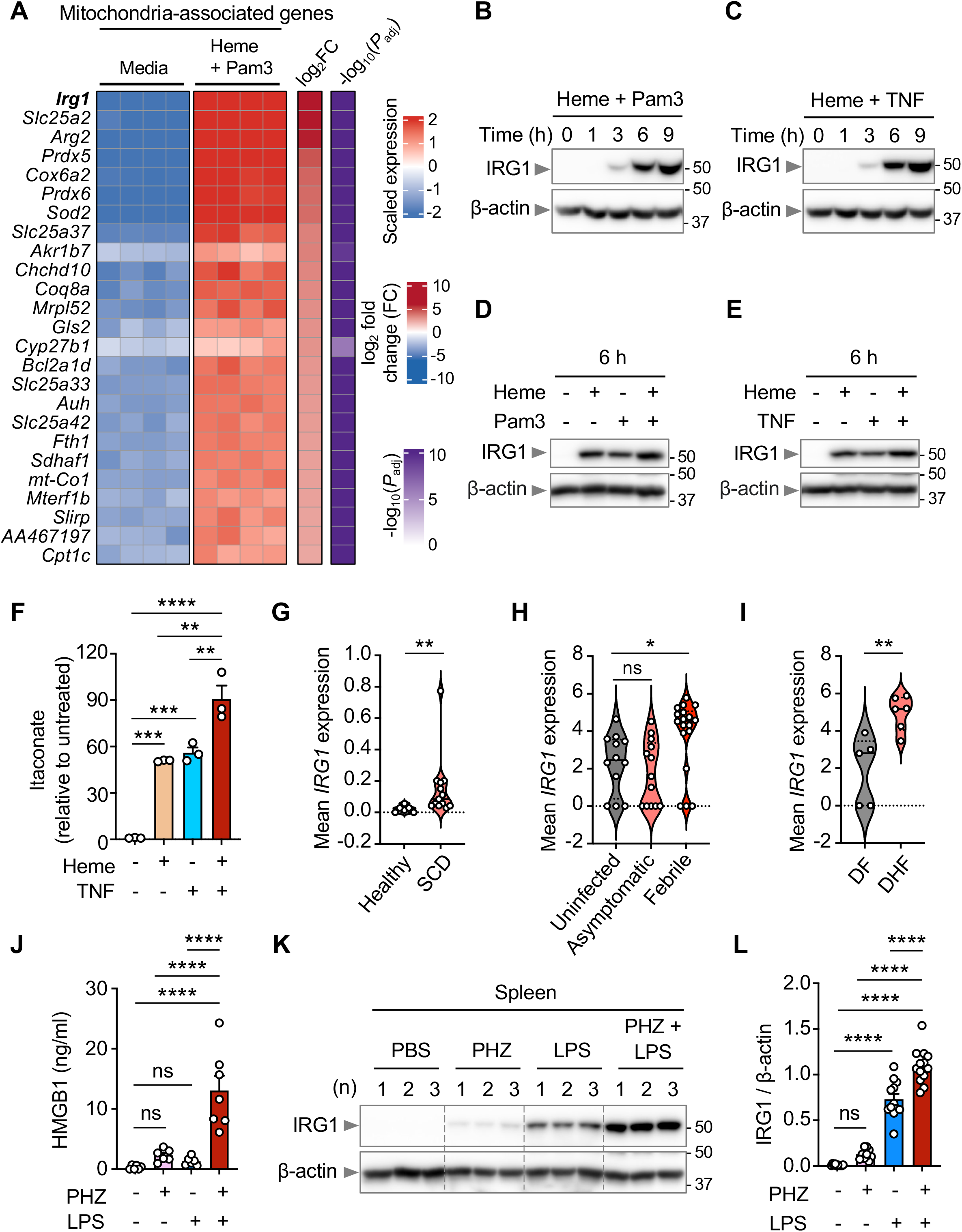
IRG1 is upregulated under hemolytic conditions. (A) Heatmap showing top differentially expressed mitochondria-associated genes in wild-type (WT) bone marrow-derived macrophages (BMDMs) stimulated with heme plus Pam3CSK4 (Pam3) for 30h, compared to media control. (B and C) Western blot analysis of immune responsive gene 1 (IRG1) expression in WT BMDMs treated with heme plus Pam3 (B) and heme plus TNF (C) at the indicated timepoints. β-actin was used as a loading control. (D and E) Western blot analysis of IRG1 expression in WT BMDMs 6 h after treatment with heme and Pam3, alone or in combination (D) or heme and TNF, alone or in combination (E). β-actin was used as a loading control. (F) Intracellular levels of itaconate in WT BMDMs 24 h after treatment with heme and TNF, alone or in combination, relative to untreated (media only) controls. (G–I) Violin plots showing mean *IRG1* expression in patients with sickle cell disease (SCD; G), malaria (H), and dengue hemorrhagic fever (DHF; I) compared with respective controls. For dengue, *IRG1* expression was compared between patients with hemorrhagic fever (DHF) and those with non-hemorrhagic dengue fever (DF). Data are from classical monocytes (CD14⁺CD16⁻) in SCD (GSE168532), and peripheral blood mononuclear cells (PBMCs) in malaria (GSE241467) and dengue (GSE215835). (J) Serum HMGB1 levels in WT mice 26 h after treatment with phenylhydrazine (PHZ, n = 6) and lipopolysaccharide (LPS, n = 6), alone or in combination (PHZ plus LPS, n = 7). Mice injected with phosphate-buffered saline (PBS) were used as control (n = 6). Pooled results from two independent experiments is shown. (K and L) Western blot analysis (K) and corresponding densitometric quantification of IRG1 expression (L) in spleen tissue lysates from WT mice 26 h after treatment with PHZ (n = 12) and LPS (n = 12), alone or in combination (PHZ plus LPS, n = 13). Mice treated with PBS served as a control (n = 10). β-actin was used as a loading control. Representative western blot, with each lane corresponding to lysate from an individual mouse, is shown in K, and pooled quantification from three independent experiments is shown in L. Data are representative of one (A), two (J) or at least three (B–F, K, and L) independent experiments. Mean ± SEM are shown (F, J, and L). Statistical analyses were performed using two-way ANOVA (F, J, and L), Mann–Whitney test (G and I), or Kruskal–Wallis test (H). \**P* < 0.05; \*\**P* < 0.01; \*\*\**P* < 0.001; \*\*\*\**P* < 0.0001; ns, non-significant.

We next sought to extend these findings to human disease and understand the expression of *IRG1*. To this end, we re-analyzed public datasets from patients with disease conditions associated with hemolysis. *IRG1* was significantly elevated in CD14^+^CD16^-^monocytes from patients with sickle cell disease (SCD)^21^ (Figure 1G) and in peripheral blood mononuclear cells (PBMCs) from patients with febrile malaria^22^ (Figure 1H). Similarly, higher *IRG1* expression was observed in PBMCs from patients with hemorrhagic dengue fever compared to non-hemorrhagic dengue fever cases^23^ (Figure 1I). Elevated *IRG1* expression was also observed in patients with familial hemophagocytic lymphohistiocytosis (fHLH)^24^ (Figure S1F), severe COVID-19 (Figure S1G), and Ebola infection (Figure S1H), as well as in macaques infected with Marburg virus (Figure S1I), all conditions commonly associated with hemolysis^25–28^. Together, these findings suggest that circulating heme may drive *IRG1* induction.

To experimentally investigate whether circulating heme contributes to IRG1 upregulation, we used a murine model mimicking hemolytic pathology. Mice were injected with the hemolysis inducer phenylhydrazine (PHZ)^29^, a sublethal dose of lipopolysaccharide (LPS), or the combination of PHZ plus LPS. PHZ plus LPS-treated mice showed significantly higher serum HMGB1 levels compared to individual treatment groups (Figure 1J), with a similar pattern observed for serum LDH levels (Figure S1J), suggesting lytic cell death was occurring in these mice. Furthermore, caspase-8 cleavage was detected only in mice treated with PHZ plus LPS, but not with individual treatments (Figure S1K), indicating that hemolysis potentiates cell death *in vivo*. We then assessed IRG1 expression in spleen tissues, where PHZ treatment caused a modest, non-significant increase in IRG1 expression, whereas LPS alone induced significant IRG1 upregulation (Figure 1K and 1L). The combination of PHZ and LPS resulted in a further increase in IRG1 expression that was significantly higher than LPS treatment alone (Figure 1K and 1L). Together with the elevated IRG1 expression observed across human diseases associated with hemolysis, these findings suggest that IRG1 may contribute to the pathogenesis of diseases with hemolysis, including infections, hemolytic diseases, and inflammatory diseases.

### IRG1 mediates inflammatory cell death, PANoptosis, through itaconate

The combination of heme with TNF or PAMPs induces robust inflammatory cell death, PANoptosis^15,16^. To determine the role of IRG1 in this inflammatory cell death, we used multiple genetic approaches. Both genetic deletion of *Irg1* (Figure S2A) and siRNA-mediated knockdown of *Irg1* (Figure S2B) significantly reduced lytic cell death in response to heme plus TNF, heme plus Pam3, and heme plus LPS (Figure 2A, 2B, S2C–F). These findings were corroborated using BMDMs from littermate mice, where *Irg1*^−/−^ BMDMs exhibited marked protection against cell death induced by heme plus TNF compared with their *Irg1*^+/+^ littermates (Figure S2G). Our RNAseq analysis had also identified several other mitochondria-associated genes that were upregulated in response to hemolytic triggers (Figure 1A and S1C). However, knockdown of these other top upregulated genes, *Slc25a2*, *Arg2*, and *Prdx5*, did not reduce cell death in response to heme plus TNF or heme plus Pam3 (Figure S2H–L). These results further support the specific role of IRG1 in regulating heme-induced cell death responses. We next assessed the role of IRG1 in human macrophages by performing siRNA-mediated knockdown of *IRG1* in differentiated THP-1 macrophages and primary human monocyte-derived macrophages (Figure S2M and S2N). IRG1 knockdown significantly reduced heme plus LPS-induced lytic cell death in both human macrophage models (Figure S2O and S2P), demonstrating that the IRG1-dependent regulation of lytic cell death is conserved in both murine and human macrophages.

**Figure 2.**
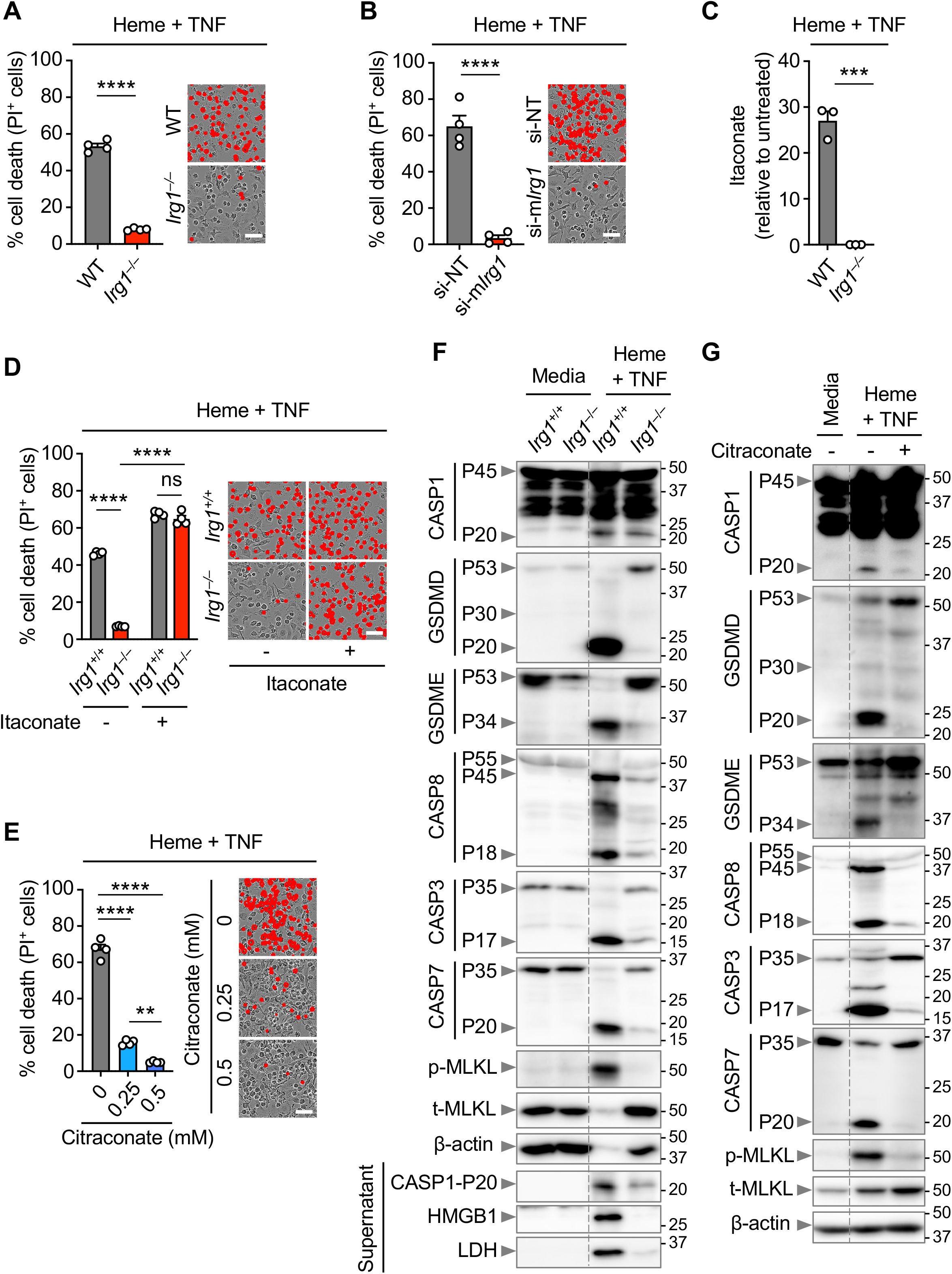
IRG1 – Itaconate axis regulates NLRC5-mediated PANoptosis. (A) Quantification showing the percentage of cell death (propidium iodide (PI)^+^ cells) and representative images of wild-type (WT) and *Irg1^−/−^* bone marrow-derived macrophages (BMDMs) treated with heme plus tumor necrosis factor (TNF) at 60 h post-treatment. (B) Quantification showing the percentage of cell death (PI^+^ cells) and representative images of WT BMDMs electroporated with control (non-targeting, NT) siRNA or mouse *Irg1*–targeting siRNA (si-m*Irg1*), treated with heme plus TNF at 48 h post-treatment. (C) Intracellular levels of itaconate in WT and *Irg1^−/−^* BMDMs 28 h after stimulation with heme plus TNF, relative to untreated (media only) controls. (D) Quantification showing the percentage of cell death (PI^+^ cells) and representative images of *Irg1^+/+^* and *Irg1^−/−^* BMDMs stimulated with heme plus TNF in the presence or absence of itaconate (5 mM) at 60 h post-treatment. Cells treated with vehicle (phosphate-buffered saline, PBS) served as a control. (E) Quantification showing the percentage of cell death (PI^+^ cells) and representative images of WT BMDMs stimulated with heme plus TNF in the presence or absence of the indicated concentrations of citraconate for 48 h. (F and G) Western blot analysis of pro- and activated caspase-1 (CASP1; P45 and P20, respectively); pro-, activated, and inactivated gasdermin D (GSDMD; P53, P30, and P20, respectively); pro- and activated gasdermin E (GSDME; P53 and P34, respectively); pro- and cleaved caspase-8 (CASP8; P55 and P45/P18, respectively); pro- and cleaved caspase-3 (CASP3; P35 and P17, respectively); pro- and cleaved caspase-7 (CASP7; P35 and P20, respectively); phosphorylated and total MLKL (p-MLKL and t-MLKL) in *Irg1*^+/+^ and *Irg1^−/−^* BMDMs stimulated with heme plus TNF for 0 h (Media) and 60 h (F), and in WT BMDMs treated with heme plus TNF with or without 0.5 mM citraconate for 0 h (Media) and 48 h (G). For loading control, β-actin is shown. In panel (F), cleaved caspase-1 (P20), HMGB1, and LDH were also measured in the culture supernatants. Scale bar = 50 µm (A, B, D, and E). Data are representative of at least three (A–G) independent experiments. Mean ± SEM are shown (A–E). Statistical analyses were performed using an unpaired t test (A–C), two-way ANOVA (D), or one-way ANOVA (E). \*\**P* < 0.01; \*\*\**P* < 0.001; \*\*\*\**P* < 0.0001; ns, non-significant.

IRG1 catalyzes the production of itaconate from the TCA cycle^20^. Itaconate has diverse and context-dependent effects across a range of diseases, from infections to cancer and neurodegeneration. For example, itaconate can reduce inflammation and tissue damage, as seen in sepsis^30^ and *Mycobacterium tuberculosis* infection^31^; in contrast, limited studies have suggested itaconate can induce inflammatory responses, such as in tissue-resident alveolar macrophages to promote acute lung injury^32^. We therefore sought to understand itaconate’s role in inflammatory cell death. We found that heme plus TNF treatment induced itaconate production in WT BMDMs, yet not in IRG1-deficient BMDMs (Figure 2C). To determine whether itaconate production directly contributed to cell death, we supplemented exogenous itaconate in *Irg1*^−/−^ BMDMs and found that this restored the cell death phenotype (Figure 2D), suggesting that IRG1 drove cell death through itaconate. These findings were further supported with pharmacological inhibition of IRG1 using a naturally occurring competitive inhibitor, citraconate^33^. Treatment with citraconate caused a dose-dependent reduction in cell death induced by heme plus TNF (Figure 2E) and heme plus Pam3 (Figure S2Q). Collectively, these results suggest that IRG1-dependent itaconate production is required for heme combination-induced cell death.

Heme in combination with TNF or PAMPs is a potent inducer of NLRC5-mediated PANoptosis^15^. We therefore sought to biochemically characterize whether IRG1 contributes to the activation of PANoptosis molecules induced by heme plus TNF treatment. Deletion of IRG1 markedly reduced the activation of PANoptotic molecules, including caspase-1, -8, -3 and -7, as well as the cleavage of gasdermin D (GSDMD, inactive P20 fragment) and gasdermin E (GSDME, active P34 fragment), along with phosphorylation of mixed lineage kinase domain-like pseudokinase (MLKL) following heme plus TNF treatment (Figure 2F). Additionally, *Irg1*^−/−^ BMDMs showed reduced HMGB1 and LDH release into the supernatant, indicating impaired plasma membrane rupture in PANoptosis (Figure 2F). A comparable protective effect was observed when IRG1 was inhibited using citraconate (Figure 2G). Together, these findings suggest that IRG1, through its enzymatic function, regulates the activation of NLRC5-mediated PANoptosis to drive inflammatory cell death.

To determine the specificity of IRG1 in cell death, we assessed the role of IRG1 across other cell death pathways and in response to other PANoptotic triggers. First, *Irg1*^+/+^ and *Irg1*^−/−^ BMDMs underwent similar cell death (Figure S3A) and ATP depletion (Figure S3B) in response to the apoptotic trigger, low dose staurosporine (STS), suggesting that IRG1 does not regulate apoptosis. The lack of PI uptake further confirmed that this trigger induced the non-lytic pathway, apoptosis (Figure S3C). Similarly, *Irg1*^+/+^ and *Irg1*^−/−^ BMDMs underwent similar cell death in response to the necroptotic trigger, TNF plus zVAD (Figure S3D) and the pyroptotic trigger, poly(dA:dT) transfection (Figure S3E). We next assessed the role of IRG1 in response to other PANoptotic triggers and found that *Irg1*^−/−^ BMDMs were partially protected from cell death induced by TNF plus IFN-γ (Figure S3F); however, no protection was observed with other PANoptotic triggers, including LPS plus ATP, KPT-330 plus IFN-β, influenza A virus (IAV) infection, or TAK1 inhibition (Figure S3G–J). These findings indicate that IRG1 regulates cell death in a context- and stimulus-specific manner.

### Glutaminolysis feeds itaconate synthesis to drive inflammatory cell death

Given that IRG1 has a key role in NLRC5-mediated PANoptosis (Figure 2A, 2B, S2C–G), and that itaconate synthesized by IRG1 is the downstream effector in this process (Figure 2D), we next investigated the upstream metabolic pathways that contribute to itaconate production and cell death. Itaconate is derived from the TCA cycle (Figure S4A), and we examined the upstream metabolic pathways that feed into the TCA cycle. We first assessed the status of glycolysis by measuring pyruvate and lactate levels (Figure S4A). Heme plus TNF treatment increased intracellular pyruvate (Figure S4B) and lactate (Figure S4C) levels, indicating enhanced glycolysis. To understand whether glycolysis contributed to NLRC5-mediated PANoptosis, we treated WT BMDMs with heme plus TNF in the presence or absence of glucose. Glucose deprivation did not impact the amount of cell death induced by heme plus TNF (Figure S4D). As an alternative approach to assess the contribution of glycolysis, we replaced glucose with galactose, which substantially limits glycolytic flux while preserving mitochondrial oxidative metabolism^34^. Heme plus TNF induced comparable levels of cell death in glucose- and galactose-containing media (Figure S4D). Together, these findings indicate that although glycolytic metabolism is increased following heme plus TNF treatment, enhanced glycolysis does not drive NLRC5-mediated PANoptosis.

Itaconate is a known inhibitor of succinate dehydrogenase (SDH; Figure S4A)^35,36^, and previous studies have demonstrated that SDH deficiency can trigger metabolic rewiring to replenish TCA intermediates^37,38^. Glutamine is one of the primary sources for replenishing TCA intermediates, particularly through α-ketoglutarate^39^ (Figure S4A). Given this, we next investigated the status of glutaminolysis following heme plus TNF stimulation. Heme plus TNF treatment decreased intracellular glutamine abundance (Figure S4E) and resulted in a significantly higher glutamate to glutamine ratio (Figure 3A) when compared to untreated BMDMs. Additionally, heme plus TNF treatment significantly increased intracellular α-ketoglutarate content (Figure S4F) and ammonia release (a byproduct of glutamine metabolism) (Figure S4G) when compared to untreated cells. Together, these results indicate that heme plus TNF stimulation enhances glutaminolysis, likely as a compensatory response to SDH inhibition by itaconate.

**Figure 3.**
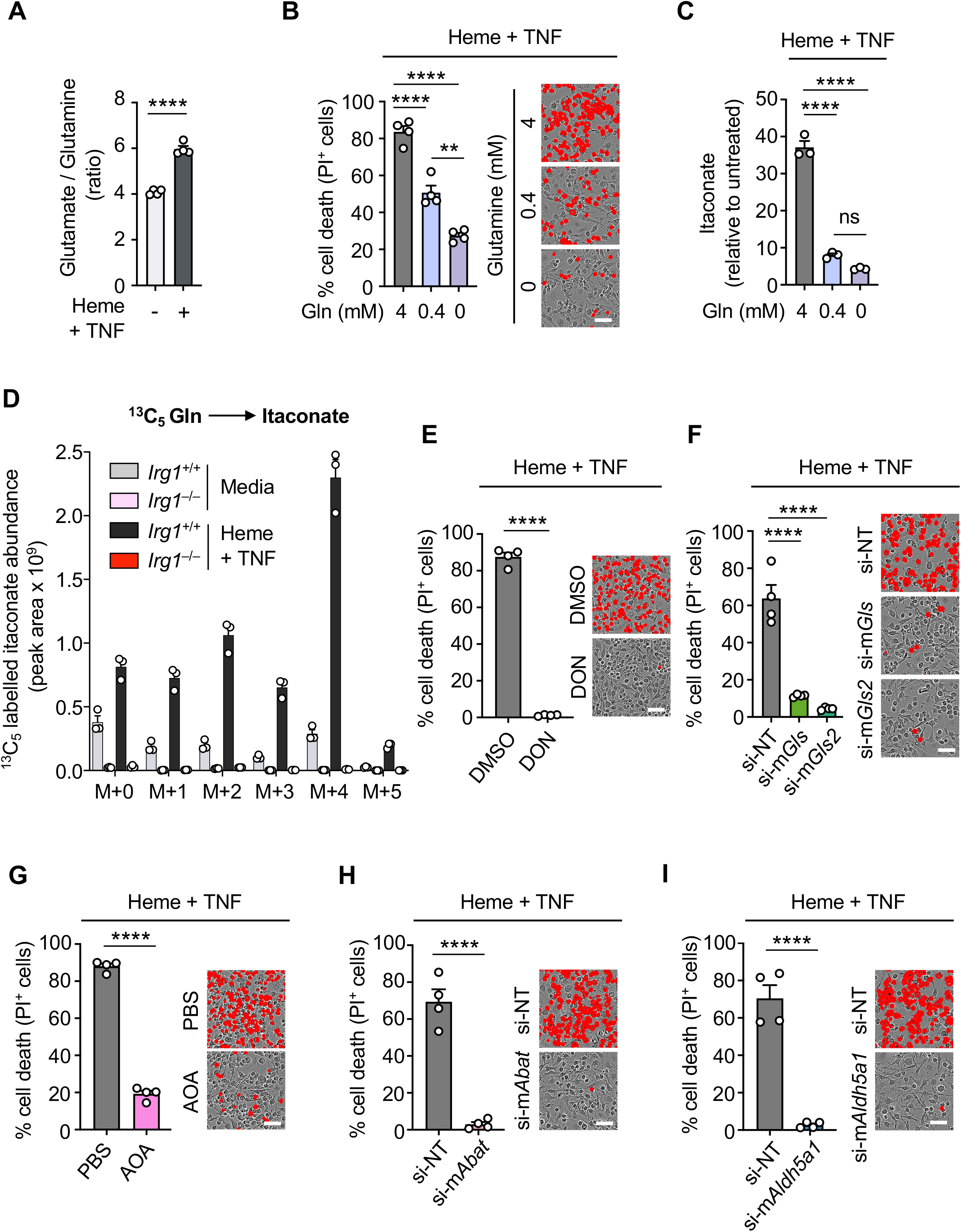
Glutamine feeds the TCA cycle through anaplerosis and GABA shunt to drive NLRC5-mediated PANoptosis. (A) Intracellular glutamate-to-glutamine ratio in wild-type (WT) bone marrow-derived macrophages (BMDMs) stimulated with or without heme plus tumor necrosis factor (TNF) for 24 h. (B) Quantification showing the percentage of cell death (propidium iodide (PI)^+^ cells) and representative images of WT BMDMs stimulated with heme plus TNF in glutamine (Gln)-free media or media containing 0.4 or 4 mM Gln for 48 h. (C) Intracellular levels of itaconate relative to untreated (media only) controls at 24 h post-treatment in response to heme plus TNF in Gln-free media or media containing 0.4 or 4 mM Gln. (D) Abundance of ^13^C_5_-labelled itaconate in *Irg1*^+/+^ and *Irg1^−/−^* BMDMs under media control conditions or in response to heme plus TNF treatment in presence of ^13^C_5_-L-glutamine (4 mM) at 16 h (E) Quantification showing the percentage of cell death (PI^+^ cells) and representative images of WT BMDMs stimulated with heme plus TNF in the presence or absence of 6-diazo-5-oxo-L-norleucine (DON, 10 µM) for 48 h. Cells were treated with vehicle (dimethyl sulfoxide, DMSO) as a control. (F) Quantification showing the percentage of cell death (PI^+^ cells) and representative images of WT BMDMs electroporated with control (non-targeting, NT) siRNA or siRNA targeting mouse *Gls* (si-m*Gls*) or *Gls2* (si-m*Gls2*) and treated with heme plus TNF for 48 h. (G) Quantification showing the percentage of cell death (PI^+^ cells) and representative images of WT BMDMs stimulated with heme plus TNF in the presence or absence of aminooxyacetic acid (AOA, 2.5 mM) for 48 h. Cells were treated with vehicle (phosphate-buffered saline, PBS) as a control. (H) Quantification showing the percentage of cell death (PI^+^ cells) and representative images of WT BMDMs electroporated with control (NT) siRNA or mouse *Abat*-targeting siRNA (si-m*Abat*) and treated with heme plus TNF for 48 h. (I) Quantification showing the percentage of cell death (PI^+^ cells) and representative images of WT BMDMs electroporated with control (NT) siRNA or mouse *Aldh5a1*-targeting siRNA (si-m*Aldh5a1*) and treated with heme plus TNF for 48 h. Scale bar = 50 µm (B, E–I). Data are representative of one (D), two (F), or at least three independent experiments (A–C, E, and G–I). Mean ± SEM are shown (A–I). Statistical analyses were performed using an unpaired t test (A, E, and G–I) or one-way ANOVA (B, C, and F). \*\**P* < 0.01; \*\*\*\**P* < 0.0001; ns, non-significant.

To understand the role of glutaminolysis in regulating NLRC5-mediated PANoptosis and the IRG1 pathway, we treated WT BMDMs with heme plus TNF in media containing normal (4 mM), reduced (0.4 mM), or no glutamine. Both glutamine restriction and deprivation significantly reduced heme plus TNF-induced cell death (Figure 3B). Similar results were obtained with heme plus Pam3 (Figure S5A) and heme plus LPS (Figure S5B), suggesting that glutamine metabolism plays a pivotal role in driving NLRC5-mediated PANoptosis. Given the robust increase in intracellular itaconate levels following heme plus TNF stimulation (Figure 1F), we next asked whether glutamine directly contributes to itaconate synthesis. To this end, we measured itaconate levels in WT BMDMs treated with heme plus TNF in normal (4 mM), reduced (0.4 mM), or glutamine-free media. Decreasing glutamine availability significantly reduced itaconate content (Figure 3C), suggesting that glutamine is used for itaconate synthesis in cells treated with heme plus TNF. To directly determine whether glutamine-derived carbon contributes to itaconate synthesis, we performed ^13^C_5_-glutamine carbon tracing. We found that ^13^C_5_-labelled itaconate accumulated in *Irg1*^+/+^ BMDMs, but not in *Irg1*^−/−^ BMDMs, in response to heme plus TNF treatment (Figure 3D). These findings align with earlier studies showing that, under inflammatory conditions, glutamine is utilized as a carbon source for itaconate synthesis^40^. Additionally, exogenous addition of itaconate completely restored cell death in glutamine deprived *Irg1*^+/+^ and *Irg1*^−/−^ BMDMs (Figure S5C), further supporting that itaconate synthesis downstream of glutaminolysis is required for cell death in response to hemolytic stimuli. Taken together, these results suggest that hemolytic triggers rewire the TCA cycle, driving glutaminolysis to fuel IRG1-dependent itaconate synthesis that leads to NLRC5-mediated PANoptosis.

### α-Ketoglutarate anaplerosis and GABA shunt pathways drive IRG1-mediated cell death

Having established that glutaminolysis contributed to itaconate synthesis and cell death in response to heme combinations (Figure 3B–D, S5A, and S5B), we next sought to further dissect the contribution of key steps in the glutaminolysis pathway that regulated IRG1-mediated cell death. Our data from glutamine-restriction and ^13^C_5_-glutamine carbon tracing experiments indicated that glutamine levels are the key determinant of intracellular itaconate accumulation upon heme plus TNF treatment (Figure 3C and 3D). Thus, we sought to determine the sources of glutamine-derived carbon entering the TCA cycle to support itaconate production and eventual cell death. Glutamine can feed into the TCA cycle through both the anaplerosis (via glutamate to α-ketoglutarate conversion) and GABA (gamma-aminobutyric acid) shunt pathways (Figure S4A). We first used DON (6-diazo-5-oxo-L-norleucine) to inhibit the glutaminases GLS and GLS2, which convert glutamine to glutamate and fuel the anaplerosis pathway (Figure S4A). DON treatment led to significant reduction in itaconate levels (Figure S5D) and cell death in response to heme plus TNF (Figure 3E). A similar protective effect was observed with heme plus Pam3 (Figure S5E). Exogenous itaconate restored heme plus TNF-induced cell death in DON treated *Irg1*^+/+^ and *Irg1*^−/−^ BMDMs (Figure S5F), supporting that itaconate is a critical cell death mediator downstream of glutaminolysis. We next genetically validated these findings by silencing *Gls* and *Gls2* using siRNAs (Figure S5G and S5H). Knockdown of either gene significantly reduced cell death induced by heme plus TNF (Figure 3F) and heme plus Pam3 (Figure S5I), confirming the contribution of glutaminase-dependent glutamine metabolism to inflammatory cell death. We then examined the next step in this pathway, the conversion of glutamate to α-ketoglutarate (the TCA metabolite), by inhibiting the transaminase enzymes (glutamate-oxaloacetate transaminase (GOT) and glutamate-pyruvate transaminase (GPT)) using aminooxyacetic acid (AOA) (Figure S4A). Addition of AOA significantly reduced itaconate levels (Figure S5J) and the cell death induced by heme plus TNF (Figure 3G). Similarly, AOA reduced heme plus Pam3-induced cell death (Figure S5K). Exogenous itaconate restored heme plus TNF-induced cell death following AOA treatment (Figure S5L), further supporting itaconate synthesis as a critical downstream step linking glutaminolysis to inflammatory cell death.

We next asked whether glutamine-derived α-ketoglutarate could also contribute to itaconate production through reductive carboxylation. Mitochondrial IDH2 catalyzes the reductive conversion of α-ketoglutarate to isocitrate, which can subsequently contribute to citrate and aconitate pools that serve as precursors for IRG1-dependent itaconate synthesis (Figure S4A). Previous studies have implicated IDH2-mediated reductive carboxylation in glutamine-derived itaconate production^41^. However, *Idh2* silencing (Figure S6A) only partially reduced the cell death in response to heme plus TNF (Figure S6B), suggesting that IDH2-dependent reductive carboxylation has a minor contribution to cell death in response to hemolytic stimuli.

In addition to anaplerosis through α-ketoglutarate, glutamine can also enter the TCA cycle through the GABA shunt pathway (Figure S4A). Using siRNA, we silenced *Abat*, which encodes the GABA-transaminase (also known as ABAT) that is responsible for converting GABA to succinic semialdehyde (SSAL; Figure S4A), eventually leading to succinate. *Abat* knockdown (Figure S6C) significantly reduced cell death in response to heme plus TNF (Figure 3H) and heme plus Pam3 (Figure S6D) when compared to non-targeting siRNA transfected cells. Similarly, silencing *Aldh5a1* (Figure S6E), which encodes succinic semialdehyde dehydrogenase (SSADH) and catalyzes the conversion of SSAL to succinate (Figure S4A), significantly reduced cell death induced by heme plus TNF (Figure 3I) and heme plus Pam3 (Figure S6F). Together, these findings indicate that the GABA shunt provides an additional route through which glutamine metabolism contributes to inflammatory cell death. Altogether, these findings indicate that glutaminolysis promotes NLRC5-mediated PANoptosis through multiple metabolic routes, including α-ketoglutarate-mediated oxidative metabolism and the GABA shunt pathway. IDH2-dependent reductive carboxylation also contributes to the phenotype, although only partially. These pathways may therefore provide multiple sources of TCA intermediates that support IRG1-dependent itaconate production and the subsequent cell death.

### Itaconate links glutaminolysis to mitochondrial ROS generation and cell death

Given that mitochondrial function is closely linked to the TCA cycle and metabolic remodeling^19^, we hypothesized that mitochondrial ROS may act as a critical bridge between the metabolic rewiring and the downstream cell death. We have previously demonstrated that ROS plays a critical role in driving NLRC5-mediated PANoptosis^15,16^. Moreover, IRG1 has been implicated in ROS generation in response to various stimuli^42,43^. Based on these observations, we hypothesized that IRG1 promotes mitochondrial ROS production in NLRC5-mediated PANoptosis. To experimentally test this, we measured mitochondrial ROS production using the MitoSOX probe. *Irg1*^−/−^ BMDMs showed significantly reduced mitochondrial ROS (mtROS) production compared to *Irg1*^+/+^ cells upon treatment with heme plus TNF (Figure 4A and 4B) and heme plus Pam3 (Figure S7A and S7B). Similar results were obtained from cells transfected with siRNA targeting *Irg1* (Figure S7C and S7D). Exogenous itaconate restored mtROS production in *Irg1*^−/−^ BMDMs (Figure 4A and 4B), suggesting itaconate is the downstream mediator of IRG1-dependent mtROS production. Moreover, exogenous itaconate further increased mtROS production in *Irg1*^+/+^ cells (Figure 4A and 4B), further implicating itaconate in mtROS generation. Consistent with a role for mtROS in promoting cell death, pharmacological attenuation of ROS with N-acetyl-L-cysteine (NAC) (Figure S7E) reduced cell death in response to heme plus TNF (Figure S7F). These data support a role for IRG1 in mediating mtROS generation, a known contributor to NLRC5-mediated PANoptosis^15,16^.

**Figure 4.**
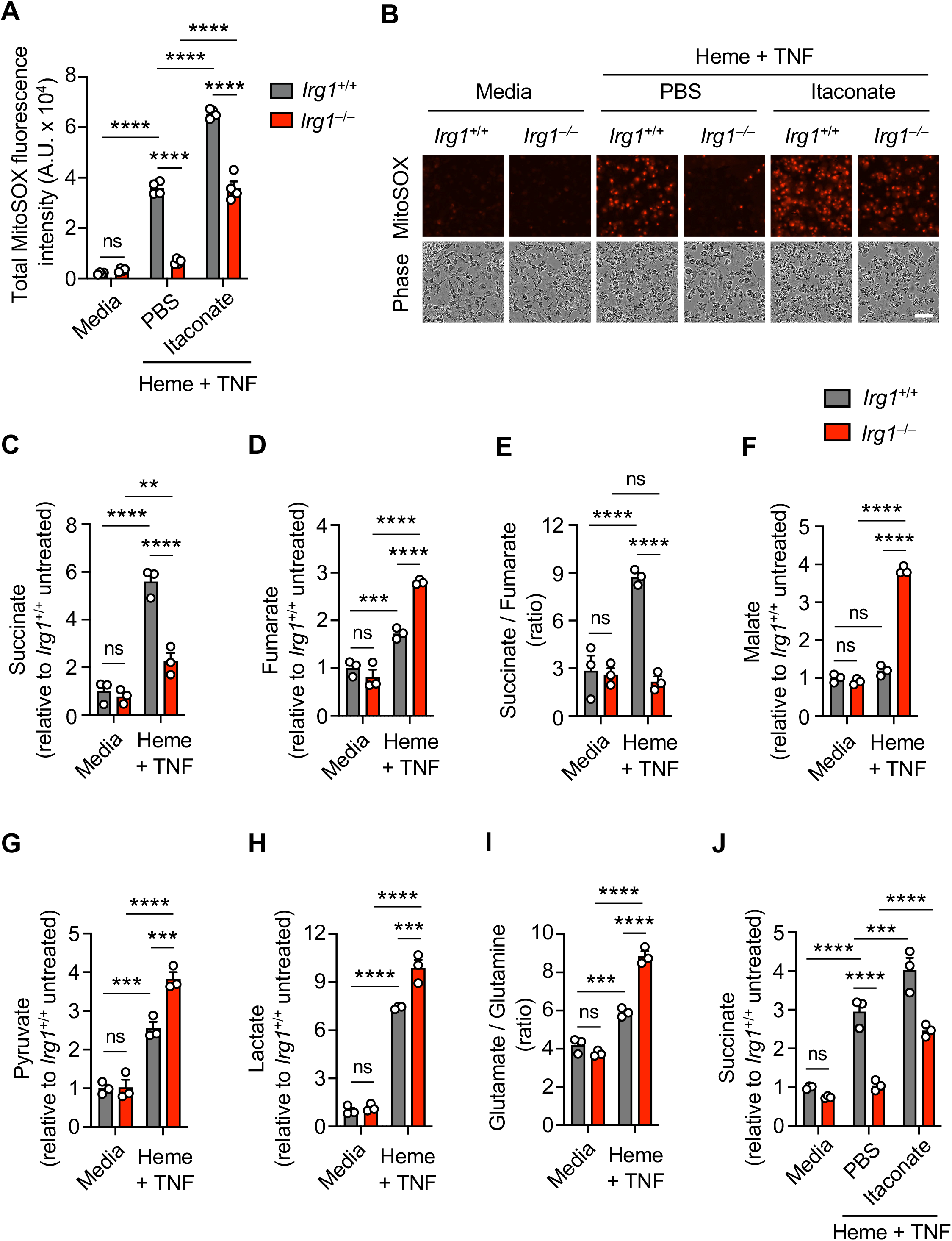
IRG1 mediates mitochondrial ROS generation and metabolic remodeling in response to hemolytic stimuli. (A and B) Quantification showing total MitoSOX fluorescence intensity (A) and representative images of MitoSOX red (top) and phase-contrast images (bottom) (B) in *Irg1*^+/+^ and *Irg1^−/−^* bone marrow-derived macrophages (BMDMs) under media control conditions or stimulated with heme plus tumor necrosis factor (TNF) in the presence or absence of itaconate (5 mM) for 48 h. (C–I) Intracellular levels of succinate (C), fumarate (D), succinate-to-fumarate ratio (E), malate (F), pyruvate (G), lactate (H), and glutamate-to-glutamine ratio (I) in *Irg1*^+/+^ and *Irg1^−/−^* BMDMs cultured under media control conditions or stimulated with heme plus TNF for 28 h. (J) Intracellular levels of succinate in *Irg1*^+/+^ and *Irg1^−/−^*BMDMs cultured under media control conditions or stimulated with heme plus TNF in the presence or absence of itaconate (5 mM) for 16 h. Scale bar: 50 µm (B). Data are representative of one (J), two (C–I), or at least three independent experiments (A and B). Mean ± SEM are shown (A and C–J). Statistical analyses were performed using two-way ANOVA (A and C–J). \*\**P* < 0.01; \*\*\**P* < 0.001; \*\*\*\**P* < 0.0001; ns, non-significant.

Given the crucial role we observed for glutamine metabolism in both itaconate synthesis and the resulting cell death, we next assessed whether glutaminolysis contributes to mtROS generation. Glutamine starvation, as well as inhibition of glutaminases with DON or inhibition of GOT/GPT with AOA, significantly reduced mtROS production (Figure S7G–I). Exogenous addition of itaconate blocked this reduction in mtROS (Figure S7G–I). Collectively, these results further support a role for itaconate in linking glutaminolysis to mtROS generation and the induction of cell death.

### IRG1-derived Itaconate drives metabolic remodeling in response to hemolytic stimuli

Because mtROS production is linked to mitochondrial metabolic activity in activated macrophages^44^, and IRG1-derived itaconate is known to remodel TCA metabolism and mitochondrial respiration^35,36^, we next examined whether the IRG1-dependent mtROS production was associated with alterations in TCA metabolism. We profiled TCA cycle metabolites in *Irg1*^+/+^ and *Irg1*^−/−^ BMDMs at 28 h post-treatment with heme plus TNF, to characterize the metabolic state before the onset of cell death. In *Irg1*^+/+^ BMDMs, heme plus TNF treatment resulted in a significant accumulation of succinate and an increased succinate/fumarate ratio, without a corresponding increase in malate (Figure 4C–F), indicating substantial TCA remodeling and a constraint at the succinate to fumarate conversion step. In contrast, while *Irg1*^−/−^ BMDMs also had a small increase in succinate levels following heme plus TNF treatment, succinate accumulation was significantly reduced compared with *Irg1*^+/+^ BMDMs (Figure 4C). Additionally, *Irg1*^−/−^ BMDMs exhibited significantly increased fumarate and malate levels (Figure 4D and 4F), suggesting altered distribution and utilization of downstream TCA cycle intermediates following IRG1 deletion.

We next examined whether these TCA alterations were accompanied by changes in other metabolic pathways. *Irg1*^−/−^ BMDMs exhibited higher pyruvate and lactate levels than *Irg1*^+/+^ BMDMs following heme plus TNF treatment (Figure 4G and 4H), consistent with enhanced glycolytic metabolism in the absence of IRG1^45^. In addition, the glutamate/glutamine ratio was increased following heme plus TNF treatment and was further elevated in *Irg1*^−/−^ BMDMs compared to *Irg1*^+/+^ cells (Figure 4I), suggesting increased glutamine utilization in the absence of IRG1.

We next asked whether restoration of itaconate could re-establish the characteristic TCA phenotype. We therefore examined the effect of exogenous itaconate at an earlier time point (16 h) during the initiation of cell death signaling, to assess the metabolic remodeling. Exogenous itaconate increased succinate accumulation (Figure 4J) and the succinate/fumarate ratio (Figure S7J) in *Irg1*^−/−^ BMDMs, bringing these levels closer to those seen in *Irg1*^+/+^. Together with the glutaminolysis experiments, these findings identify distinct TCA metabolic remodeling in *Irg1*^+/+^ and *Irg1*^−/−^ BMDMs in response to hemolytic stimulation and suggest that IRG1-derived itaconate couples glutamine metabolism to the metabolic and cell death responses.

### Itaconate promotes ROS and cell death through an SDH-independent mechanism in response to hemolytic stimuli

Mitochondrial ETC activity regulates electron flow, membrane potential, and cellular bioenergetics, which can influence mitochondrial ROS production and inflammatory signaling^46,47^. Given that itaconate is a competitive inhibitor of SDH that can promote succinate accumulation^35,36^, we observed increased succinate levels following heme plus TNF treatment, which were reduced in *Irg1*^−/−^ BMDMs and restored by exogenous itaconate (Figure 4C and 4J). We therefore asked whether SDH inhibition was sufficient to reproduce the mtROS and cell death effects of itaconate. However, treatment with dimethylmalonate (DMM), a competitive SDH inhibitor, did not restore mtROS (Figure 5A and 5B) or cell death (Figure 5C and 5D) in *Irg1*^−/−^ BMDMs upon heme plus TNF treatment, indicating that SDH inhibition alone is insufficient to account for the effects of itaconate in response to hemolytic stimuli.

**Figure 5.**
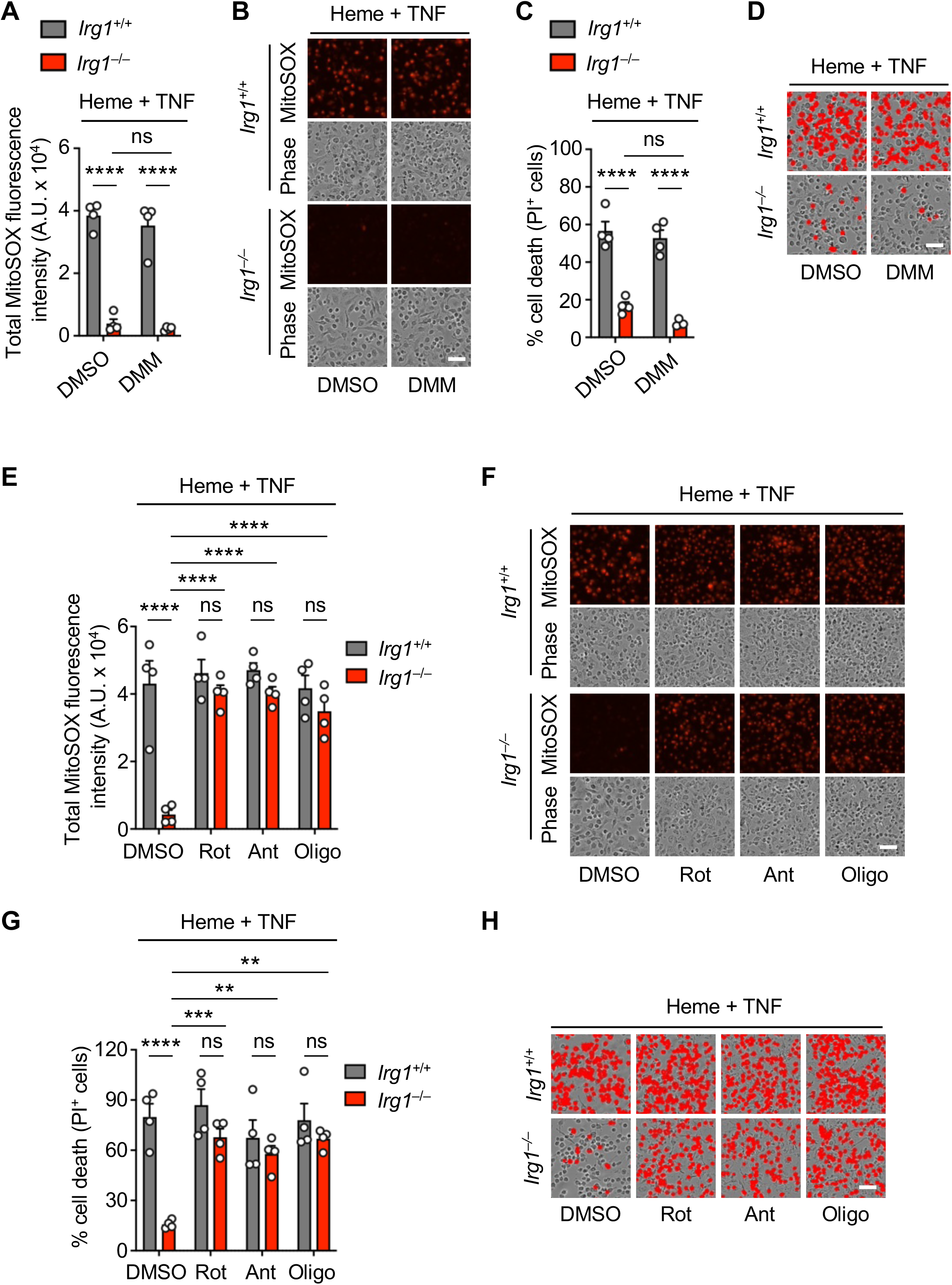
Itaconate promotes ROS and cell death through an SDH-independent mechanism in response to hemolytic stimuli. (A and B) Quantification showing total MitoSOX fluorescence intensity (A) and representative images of MitoSOX red and phase-contrast images (B) in *Irg1*^+/+^ and *Irg1^−/−^* bone marrow-derived macrophages (BMDMs) treated with heme plus tumor necrosis factor (TNF) in the presence or absence of dimethyl malonate (DMM, 5 mM) for 48 h. (C and D) Quantification showing the percentage of cell death (propidium iodide (PI)^+^ cells) (C) and representative images (D) of *Irg1*^+/+^ and *Irg1^−/−^* BMDMs treated with heme plus TNF in the presence or absence of DMM for 60 h. (E and F) Quantification showing total MitoSOX fluorescence intensity (E) and representative images of MitoSOX red and phase-contrast images (F) in *Irg1*^+/+^ and *Irg1^−/−^* BMDMs treated with heme plus TNF in the presence or absence of rotenone (Rot, 6.25 nM), antimycin A (Ant, 25 nM), or oligomycin A (Oligo, 25 nM) for 48 h. (G and H) Quantification showing the percentage of cell death (PI^+^ cells) (G) and representative images (H) of *Irg1*^+/+^ and *Irg1^−/−^* BMDMs treated with heme plus TNF in the presence or absence of Rot, Ant, or Oligo for 60 h. Data in (E and F) are from an independent experiment from data in (G and H). Scale bar = 50 µm (B, D, F, and H). Data are representative of two (A, B, E, and F) or at least three independent experiments (C, D, G, and H). Mean ± SEM are shown (A, C, E, and G). Statistical analyses were performed using two-way ANOVA (A, C, E, and G). \*\**P* < 0.01; \*\*\**P* < 0.001; \*\*\*\**P* < 0.0001; ns, non-significant.

SDH links the TCA cycle to mitochondrial complex II in the ETC, where it catalyzes succinate oxidation and contributes electrons to the respiratory chain. Given that complex II inhibition using DMM was insufficient to reproduce the effects of itaconate, and previous studies have shown that IRG1-derived itaconate can alter mitochondrial bioenergetics, including mitochondrial substrate-level phosphorylation^48^, we next asked whether perturbation of mitochondrial electron transport at sites other than complex II could bypass the requirement for itaconate. We therefore examined the effects of inhibitors targeting distinct components of the mitochondrial respiratory chain, including complex I , complex III, and ATP synthase. We found that inhibition of complex I with rotenone, complex III with antimycin A, or ATP synthase with oligomycin in the context of heme plus TNF stimulation led to similar levels of mtROS production and cell death between *Irg1*^+/+^ and *Irg1*^−/−^ BMDMs (Figure 5E–H). Together, these findings indicate that perturbation of mitochondrial respiration at sites distinct from complex II can restore mtROS and cell death in the absence of IRG1, whereas SDH inhibition alone is insufficient. Consistent with our earlier finding that NAC reduced heme plus TNF-induced mtROS accumulation and cell death (Figure S7E and S7F), these data support mtROS as a convergent effector linking IRG1-derived itaconate to NLRC5-mediated PANoptosis. Thus, the effects of itaconate on mtROS and cell death extend beyond its canonical role as an SDH inhibitor.

### IRG1 promotes NLRC5-PANoptosome formation to drive PANoptosis

Given the link between mitochondrial redox state, bioenergetics and ROS production, we next assessed whether NAD^+^ depletion contributes to the IRG1-dependent cell death response. NAD^+^ is a critical cofactor maintaining mitochondrial functions and redox balance. Depletion of NAD^+^ has been linked to mitochondrial dysfunction, ROS generation, and cell death^49–52^, particularly NLRC5-mediated PANoptosis^15^. We found that NAD^+^ levels were similarly depleted upon heme plus TNF treatment in both WT and *Irg1*^−/−^ BMDMs (Figure 6A), suggesting that IRG1 likely acts downstream of the NAD^+^ depletion. Supporting this, we observed that IRG1 expression in response to heme plus TNF was significantly delayed upon supplementation with the NAD^+^ precursor nicotinamide (NAM) (Figure 6B and 6C). This mirrored a similar delay in NLRC5 expression in response to heme plus TNF upon NAM supplementation (Figure 6C), as previously demonstrated^15^. In contrast, supplementation with the antioxidant NAC had no impact on IRG1 expression in response to heme plus TNF (Figure 6C), suggesting that ROS production occurs downstream of IRG1. Together with the observed reduction in ROS in *Irg1*^−/−^ BMDMs (Figure 4A, 4B, and S7A–D), these results suggest that NAD^+^ availability may provide a metabolic link between innate immune activation, mtROS production and inflammatory cell death, and that this is in part due to the regulation of IRG1 expression.

**Figure 6.**
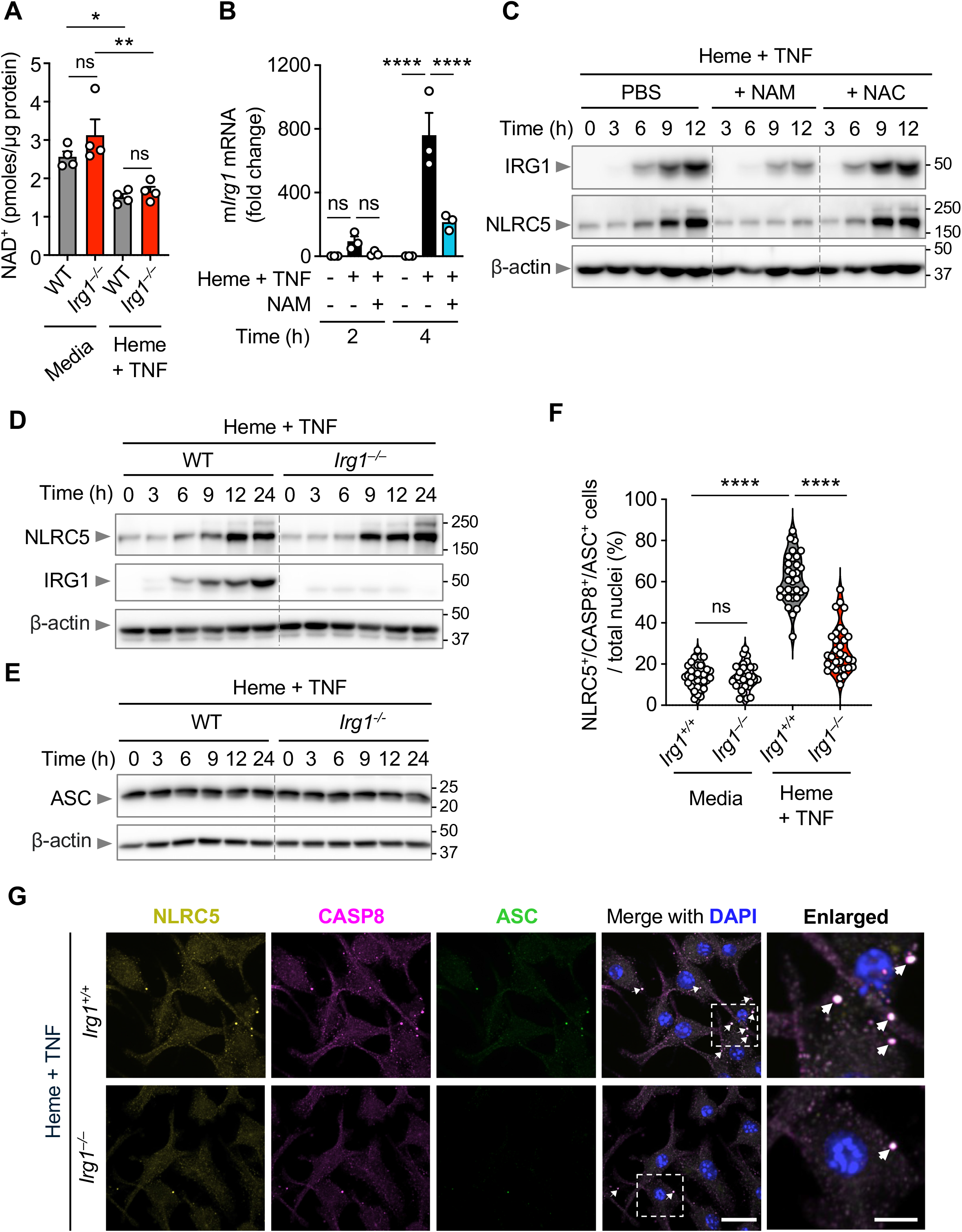
IRG1 promotes NLRC5-PANoptosome formation to drive PANoptosis. (A) Quantification of intracellular nicotinamide adenine dinucleotide (NAD^+^) levels in wild-type (WT) and *Irg1^−/−^* bone marrow-derived macrophages (BMDMs) stimulated with heme plus tumor necrosis factor (TNF) for 28 h compared to media controls. (B) *Irg1* mRNA expression at 2 and 4 h in WT BMDMs stimulated with heme plus TNF in the presence or absence of nicotinamide (NAM, 10 mM). *18S* was used for normalization of mRNA expression. Each data point represents an independent experiment (n = 3). (C) Time-course western blot analysis of IRG1 and NLRC5 expression in WT BMDMs stimulated with heme plus TNF in the presence or absence of NAM (10 mM) or N-acetyl-L-cysteine (NAC, 5 mM). β-actin was used as the loading control. (D and E) Time-course western blot analysis of NLRC5 and IRG1 expression (D) and ASC expression (E) in WT and *Irg1^−/−^* BMDMs stimulated with heme plus TNF. β-actin was used as the loading control. (F) Quantification showing the percentage of NLRC5⁺CASP8⁺ASC⁺ speck-containing cells among total nucleated cells in *Irg1*^+/+^ and *Irg1^−/−^* BMDMs at 46 h post-stimulation with heme plus TNF or media only as a control. Each data point represents a single field of view. Pooled quantification from three independent experiments is shown. (G) Representative images of *Irg1*^+/+^ and *Irg1^−/−^*BMDMs stimulated with heme plus TNF for 46 h and stained for NLRC5, caspase-8 (CASP8), and ASC, with DAPI counterstaining for nuclei. The boxed area (merged) is shown at higher magnification on the right (enlarged). White arrows indicate NLRC5^+^ CASP8^+^ ASC^+^ colocalized specks. Scale bars: 15 μm (G, merge column), and 7 μm (G, enlarged column). Data are representative of at least three independent experiments (A–G). Mean ± SEM are shown (A and B). Statistical analyses were performed using two-way ANOVA (A, B, and F). \**P* < 0.05; \*\**P* < 0.01; \*\*\*\**P* < 0.0001; ns, non-significant.

Given that NLRC5 and IRG1 were both downstream of NAD^+^ (Figure 6B and 6C) and that IRG1 was necessary for induction of PANoptosis (Figure 2F and 2G), we next assessed whether IRG1 regulates NLRC5 expression. NLRC5 levels were comparable between WT and *Irg1*^−/−^ BMDMs upon heme plus TNF treatment (Figure 6D), indicating that IRG1 does not influence NLRC5 protein expression. Similarly, ASC expression was comparable between WT and *Irg1*^−/−^ BMDMs (Figure 6E), suggesting that IRG1 does not broadly regulate the expression of PANoptosome components. Downstream of its expression, NLRC5 forms a PANoptosome complex to mediate inflammatory cell death, PANoptosis^15^. Therefore, we next assessed whether NLRC5-PANoptosome assembly was influenced by IRG1. We observed a significant decrease in NLRC5 colocalizing with other PANoptosome complex members, caspase-8 and ASC, in *Irg1*^−/−^ BMDMs (Figure 6F and 6G), indicating that IRG1 mediates NLRC5-PANoptsome assembly to drive cell death. Together, these findings suggest that NAD^+^ depletion triggers IRG1 expression, leading to mtROS generation and NLRC5-PANoptosome assembly to promote cell death.

## DISCUSSION

Metabolic control of immune signaling and cell death is increasingly recognized as a key factor in many diseases, including cancer, neurodegenerative diseases, autoimmune disorders, infections and sepsis. Beyond serving as metabolic intermediates, specific metabolites can function as signaling molecules affecting cytokine release, redox balance, and cell survival^4–9^. Here, we show that IRG1 drives itaconate synthesis in response to hemolytic triggers, leading to mitochondrial ROS production and NLRC5-dependent inflammatory cell death, PANoptosis. Our findings further identify a metabolic pathway linking NAD^+^ availability and glutamine metabolism to IRG1-dependent itaconate production, mitochondrial metabolic remodeling, ROS generation, and inflammatory cell death.

Our results demonstrate a pro-inflammatory and pro-death role for IRG1. Previous studies have primarily defined anti-inflammatory roles for IRG1 and its metabolite itaconate^53^ in limiting cytokine release and oxidative stress during sepsis and infections^30,31,54^, as well as in models of ischemic liver and brain injury^55–57^. Here, we found that hemolytic conditions promoted glutaminolysis, rewiring the TCA cycle to fuel itaconate synthesis that led to inflammatory lytic cell death, PANoptosis. Glutamine-derived carbon contributed to this pathway through multiple routes, including canonical α-ketoglutarate metabolism and the GABA shunt, highlighting the integration of glutamine metabolism to itaconate production and ensuing cell death.

Similarly, other recent studies also suggested pro-inflammatory roles for itaconate, suggesting that it can exacerbate lung injury by promoting pro-inflammatory responses in tissue-resident alveolar macrophages^32^ and that it can drive type I IFN production^53,58^. These collective findings therefore suggest that IRG1 and itaconate exert context-dependent functions, likely reflecting differences in timing of IRG1 induction or substrate availability for itaconate synthesis. Our finding that SDH inhibition alone does not recapitulate the effects of IRG1-dependent itaconate further suggests that mechanisms beyond SDH inhibition contribute to mtROS and inflammatory cell death. Consistent with this possibility, itaconate has been shown to covalently modify metabolic enzymes, including glycolytic enzymes such as GAPDH, ALDOA, and LDHA, thereby influencing cellular metabolism^45,59^. In addition, Itaconate can alkylate cysteine residues on immune sensors including NLRP3, altering their activity^60,61^. Whether similar protein modification contributes to NLRC5-PANoptosome assembly under hemolytic conditions remains to be determined.

Our findings further place NAD^+^ availability upstream of IRG1. Although NAD^+^ depletion occurs across multiple forms of cell death, restoration of NAD^+^ has been shown to preferentially suppress PANoptosis^62^, suggesting that NAD^+^ can regulate inflammatory cell death through mechanisms extending beyond IRG1. Consistent with this broader role, IRG1 deletion did not protect against several other cell-death triggers, indicating that the effects of NAD^+^ restoration are not exclusively mediated through IRG1. Thus, NAD^+^ depletion may represent a common metabolic consequence of cellular stress that, depending on the inflammatory context, engages distinct downstream pathways including IRG1.

Collectively, our study identifies IRG1 as a key metabolic regulator connecting inflammatory signaling to the enigmatic innate immune sensor NLRC5 and inflammatory cell death, PANoptosis. NAD⁺ availability regulates IRG1 expression, while glutamine metabolism supplies carbon for itaconate synthesis, providing a metabolic link between innate immune activation and inflammatory cell death. Our findings suggest that targeting this IRG1–itaconate–NLRC5 axis may offer new therapeutic opportunities to mitigate tissue injury in hemolytic, infectious, and inflammatory diseases.

## ACKNOWLEDGEMENTS

We thank all the members of the Kanneganti laboratory for their comments and suggestions during the development of this manuscript. We also thank R. Tweedell, PhD and K. Zengeler, PhD, for scientific editing and writing support. We also thank the St. Jude Hartwell Center Genome Sequencing Facility for facilitating RNA sequencing and processing. Work from our laboratory is supported by the US National Institutes of Health (AI101935, AI124346, AI160179, AR056296, and CA253095 to T.-D.K.) and the American Lebanese Syrian Associated Charities (to T.-D.K.). The content is solely the responsibility of the authors and does not necessarily represent the official views of the National Institutes of Health.

## AUTHOR CONTRIBUTIONS

N.P., B.S., and T.-D.K. conceptualized the study; N.P. and B.S. designed the methodology; N.P., B.S., A.P.Y., and R.S. performed the experiments; S.P.K., and C.A.E.F conducted the gene expression and publicly available dataset analysis; G.P. acquired and performed the mass spectrometric analysis; N.P. wrote the manuscript with input from all authors. T.-D.K. acquired the funding and provided overall supervision.

## DATA AVAILABILITY

The datasets generated during the current study are available at Gene Expression Omnibus (GEO) under accession number GSE314427, and publicly available datasets analyzed can be found in the GEO database (GSE168532, GSE241467, GSE215835, GSE26050, GSE207625, GSE83563, and GSE58287).

## METHODS

### Mice

*Irg1*^−/−^ mice (C57BL/6NJ-Acod1^em1(IMPC)J^/J; #029340) and their respective wild-type (WT) control strain (C57BL/6NJ mice; #005304) were purchased from The Jackson Laboratory. Heterozygous *Irg1*^+/−^ mice were generated in-house by crossing WT (C57BL/6NJ) and *Irg1*^−/−^ mice. *Irg1*^+/−^ mice were then intercrossed to generate littermate *Irg1*^+/+^ and *Irg1*^−/−^ mice, as well as additional *Irg1*^+/+^ controls. WT C57BL/6J (#000664; purchased from The Jackson Laboratory) and *Nlrc5^−/−^*^63^ mice were used for other experiments. All mice were bred at the Animal Resources Center at St. Jude Children’s Research Hospital and maintained under specific pathogen-free conditions. Male and female 6-to 12-week-old mice were used for in vitro experiments, and age-and sex-matched, 6-to 8-week-old male mice were used for in vivo experiments in this study. Mice were maintained with a 12 h light/dark cycle and were fed standard chow. Animal studies were conducted under protocols approved by the St. Jude Children’s Research Hospital committee on the Use and Care of Animals. Experimental animals were housed with mixed bedding for 1–2 weeks before performing in vivo experiments.

### Bone marrow-derived macrophage (BMDM) generation

Primary mouse BMDMs from WT and indicated mutant mice were grown for 6 days in IMDM (12440053, Thermo Fisher Scientific) supplemented with 10% heat-inactivated fetal bovine serum (HI-FBS; S1620, Biowest), 30% L929 conditioned media, 1% non-essential amino acids (11140-050, Thermo Fisher Scientific), and 1% penicillin and streptomycin (15070-063, Thermo Fisher Scientific). BMDMs at a density of 2.5 × 10^6^ cells/well in 6-well plates, 1 × 10^6^ cells/well in 12-well plates, or 5 × 10^5^ cells/well in 24-well plates were seeded into growth media overnight before use. For experiments involving siRNA-mediated gene knockdown, cells were resuspended in phosphate-buffered saline (PBS) at a density of 1 × 10^7^ cells/mL. For transfection by electroporation (Neon Transfection System, MPK5000, Thermo Fisher Scientific), 5 pmoles of siRNA per million cells were used. The siRNAs included non-targeting control siRNA (D-001206-14-20) and mouse-specific siRNAs targeting *Irg1* (M-048308-01-0005), *Gls* (M-043336-01-0005), *Gls2* (M-063540-01-0005) *Abat* (M-064095-00-0005), *Aldh5a1* (M-054629-01-0005), *Idh2* (M-055489-00-0005), *Slc25a2* (M-055560-01-0005), *Arg2* (M-043749-01-0005), and *Prdx5* (M-056883-01-0005), all from Horizon Discovery. Cells were used for experiments 48 h after transfection.

### Cell stimulation

Primary mouse BMDMs from WT and indicated mutant mice and siRNA-treated cells were treated in DMEM (11995-065, Gibco) supplemented with 10% HI-FBS and 1% penicillin-streptomycin. Cells were stimulated in a final volume of 250 µL, 500 µL, or 1 mL media per well when seeded in 24-, 12-, or 6-well plates, respectively. Cells were stimulated with the following DAMPs, PAMPs, cytokines, and inhibitors, either alone or in combination as indicated: 50 µM Hemin (heme; H9039, Sigma), 15 ng/mL ultrapure LPS from *E. coli* (0111: B4; tlrl-3pelps, InvivoGen), 500 ng/mL Pam3CSK4 (Pam3; tlrl-pms, InvivoGen), 100 ng/mL TNF (315-01A, Peprotech), 5 mM itaconate (I29204, Sigma), 0.25 or 0.5 mM citraconate (C82604, Sigma), 10 mM nicotinamide (NAM; N3376, Sigma-Aldrich), 5 mM N-acetyl L-cysteine (NAC; A9165, Sigma-Aldrich), 2.5 or 10 µM 6-diazo-5-oxo-L-norleucine (DON; 17580, Cayman Chemical), 1.25 or 2.5 mM aminooxyacetic acid (AOA; 28298, Cayman Chemical), 5 mM dimethyl malonate (DMM; HY-Y1787, MedChemExpress), 6.25 nM rotenone (Rot; R8875, Sigma), 25 nM antimycin A (Ant; 33357, Cell Signaling Technology [CST]), and 25 nM oligomycin A (Oligo; 4110, Tocris). For glucose deprivation experiments, BMDMs were stimulated in glucose-free DMEM (A1443001, Gibco) supplemented with 4 mM glutamine (25030081, Gibco) and 1 mM sodium pyruvate (11360070, Gibco) with or without supplementation of 25 mM glucose (15023021, Gibco) or 25 mM galactose (G5388, Sigma-Aldrich). BMDMs were also stimulated in glutamine-depleted media (10313-021, Gibco) with or without supplementation of 0.4 mM or 4 mM glutamine (25030-081, Gibco). Hemin (100 mM stock) was freshly prepared before each experiment by dissolving in filter sterilized 0.1 M NaOH, as previously described^64^. Stock solutions of DON, DMM, rotenone, antimycin A, and oligomycin A were prepared in DMSO and stored at -80°C prior to use. Stock solutions of itaconate, citraconate, NAM, NAC, and AOA were freshly prepared in respective media before use. Control cells were treated with equivalent volumes of respective vehicle solution. Unless otherwise noted in their respective methods sections, for assays comparing *Irg1*^−/−^ BMDMs with littermates or matched C57BL6/NJ WT controls, BMDMs were stimulated with 100 µM Hemin, 30 ng/mL ultrapure LPS, 1 µg/mL Pam3, or 200 ng/mL TNF, either alone or in combination.

To induce necroptosis, BMDMs were pretreated with 25 µM Z-VAD(OMe)-FMK (zVAD; 14463, Cayman Chemical) for 30 min and then stimulated with 50 ng/mL TNF (315-01A, PeproTech) for 16 h. For pharmacological inhibition of necroptosis, 30 µM Necrostatin 2 racemate (Nec-1s; S8641, Selleckchem) was added together with zVAD where indicated.

To induce AIM2-dependent pyroptosis, cells were transfected with poly(dA:dT) (tlrl-patn, InvivoGen). Each reaction consisted of 2 μg per million cells poly(dA:dT) resuspended in PBS and mixed with 0.6 μL Xfect polymer in Xfect reaction buffer (631318, Clontech Laboratories). After 10 min, DNA complexes were added to BMDMs in Opti-MEM (31985–070, Thermo Fisher Scientific). To inhibit pyroptosis, 25 µM VX-765 (HY-13205, MedChemExpress) or 20 µM suramin (574625, Sigma-Aldrich) was added added 30 min prior to transfection.

To induce PANoptosis using additional triggers, BMDMs were stimulated with 25 ng/mL TNF plus 50 ng/mL IFN-γ (315-05, PeproTech), 5 µM KPT-330 (S7252, Selleckchem) plus 50 ng/ml IFN-β (12400-1, PBL Assay), or 100 nM (5Z)-7-oxozeanol (TAK1 inhibitor [TAK1i], 17459, Cayman Chemical). For LPS plus ATP stimulation, BMDMs were primed with 100 ng/ml LPS for 4 h and then stimulated with 5 mM ATP (101275310001; Roche) for 60 min. To inhibit LPS plus ATP-induced cell death, 25 µM VX-765 or 5 µM MCC950 (inh-mcc, InvivoGen) was added 30 min prior to ATP addition. Influenza A virus (IAV; A/Puerto Rico/8/34, H1N1 [PR8]) was prepared as previously described^65^ and propagated in 11-day-old embryonated chicken eggs by allantoic inoculation. Viral titers were determined by plaque assay in Madin-Darby canine kidney (MDCK) cells (CCL-34, ATCC). For infection, BMDMs were inoculated with IAV at a multiplicity of infection (MOI) of 20 in plain DMEM (D6171, Sigma-Aldrich). Then, 10% HI-FBS was added to the cells 1 h post-inoculation.

To induce apoptosis, BMDMs were stimulated with 0.5 µM staurosporine (S5921, Sigma-Aldrich), and cell viability was assessed after 9 h.

### Human monocyte-derived macrophages

Fresh human blood was collected from the apheresis rings of anonymous healthy blood donors from the blood bank of St. Jude Children’s Research Hospital, following a protocol reviewed and approved by the St. Jude IBC. Peripheral blood mononuclear cells (PBMCs) were isolated using lymphoprep solution (#07801/07811, Stemcell Technologies). Naive monocytes were purified from the PBMCs using monocyte isolation kit (EasySep direct human monocyte isolation kit, #19669, Stemcell Technologies) according to the manufacturer’s protocol. These purified monocytes were further differentiated into monocyte-derived macrophages by culturing in RPMI media (10-040-CV, Corning) supplemented with 10% HI-FBS, 1% penicillin-streptomycin, and 25 ng/ml human M-CSF (300-25, Peprotech) for 8 days in a CO2 incubator supplied with 5% CO2 in a humidified atmosphere. Additional M-CSF-containing medium was added on days 2, 4, and 6. On day 8, adherent macrophages were harvested, washed three times with PBS, and resuspended at a cell density of 1 × 10^7^ cells/ml. Cells were transfected with 5 pmoles non-targeting (control) siRNA (D-001206-14-20, Horizon Discovery) or human specific *IRG1* siRNA (M-180668-01-0005, Horizon Discovery) per million cells using the Neon NxT Electroporation System (N10096, Invitrogen). Post-transfection, the cells were maintained in RPMI medium supplemented with 10% HI-FBS, 1% penicillin-streptomycin, and 25 ng/ml human M-CSF along with 1 ng/ml ultrapure LPS from E. coli (0111: B4) (tlrl-3pelps, InvivoGen) and 20 ng/ml human IFN-γ (300-02, Peprotech) to polarize the cells to canonical macrophages. At 48 h post-transfection, cells were washed once with PBS and treated with heme (50 µM) and LPS (15 ng/ml) in RPMI medium supplemented with 10% HI-FBS and 1% penicillin-streptomycin.

### THP-1 cells

THP-1 cells (TIB-202, ATCC) were maintained in RPMI medium supplemented with 2 mM glutamine, 10% FBS, and 1% penicillin-streptomycin. Cells were transfected with 5 pmoles siRNA (per million cells) targeting human *IRG1* or non-targeting control siRNA using the Neon NxT Electroporation System and maintained in medium containing 125 nM PMA (phorbol 12-myristate 13-acetate) to induce differentiation. After 24 h, cells were washed once with PBS and rested in RPMI for an additional 24 h. Cells were then treated with heme (100 µM) and LPS (50 ng/mL) in complete RPMI containing 4 mM glutamine.

### Real-time imaging for cell death

The kinetics of cell death were monitored using the IncuCyte S3 or SX5 (Sartorius) live-cell analysis system. BMDMs were seeded in 24-well tissue culture plates; THP-1 cells and human monocyte-derived macrophages were seeded in 48-well tissue culture plates. Cells were treated with the indicated stimuli. Cell death was measured by propidium iodide (PI; P3566, Life Technologies) incorporation following the manufacturer’s protocol. The plate was scanned for the indicated time durations, with fluorescent and phase-contrast images acquired in real-time. PI-positive dead cells are marked with a red mask and were quantified using the software package supplied with the IncuCyte imager.

### Cell viability assessment

For MTT-based cell viability measurements, the MTT Cell Proliferation Assay Kit (K2249, APExBIO) was used with minor modifications to the manufacturer’s protocol. BMDMs were seeded at 2 × 10^5^ cells/well in 48-well plates. Following indicated treatments, MTT solution was added at 10% of the total treatment volume, and cells were incubated at 37°C for 3 h. Plates were then centrifuged, and the medium was gently removed. Formazan crystals were dissolved in 150 µL DMSO, and absorbance was measured at 570 nm using a microplate reader.

As an alternative measure of cell viability, ATP levels were assessed in BMDMs seeded at 1 × 10^5^ cells/well in 96-well plates using the CellTiter-Glo Luminescent Cell Viability Assay (G7573, Promega) according to the manufacturer’s instructions.

### Detection of mitochondrial ROS

Mitochondrial ROS production was detected by staining treated BMDMs with MitoSOX red dye (M36008, Thermo Fisher Scientific), and the generation of ROS was monitored using the IncuCyte live-cell analysis system (Sartorius). Briefly, BMDMs were washed once with 1× PBS, and treated as indicated in the presence of 2.5 µM MitoSOX red dye, and the images were acquired using the IncuCyte system.

### Immunoblot analysis

After appropriate treatments, cells were lysed along with culture supernatants in caspase lysis buffer (containing 10% NP-40, 25 mM DTT and 1× protease and phosphatase inhibitors (11697498001 and 04906837001, respectively, Roche)) and 4× SDS sample loading buffer (with 2-mercaptoethanol) for probing caspase activation. For immunoblot assessment of signaling activation, culture supernatants were removed, cells were washed once with 1× Dulbecco’s PBS (DPBS; 14190-250, Thermo Fisher Scientific) and lysed in RIPA buffer and 4× SDS sample loading buffer. For analysis of caspase-1, HMGB1, and LDH release by immunoblotting, culture supernatants were collected and centrifuged at 10,000 rpm for 2 min, and the clarified, cell-free supernatants were then combined with 4× SDS sample loading buffer. All samples were boiled at 100°C for 10 min before being loaded onto gels. Proteins were resolved on 8–12% polyacrylamide gels and transferred onto PVDF membranes (IPVH00010, Millipore) using the Trans-Blot Turbo system. After blocking non-specific binding with 5% skim milk, membranes were incubated overnight with the following primary antibodies against: caspase-1 (AG-20B-0042, AdipoGen, 1:1000), caspase-3 (#9662, CST, 1:1000), cleaved caspase-3 (#9661, CST, 1:1000), caspase-7 (#9492, CST, 1:1000), cleaved caspase-7 (#9491, CST, 1:1000), caspase-8 (#4927, CST, 1:1000), cleaved caspase-8 (#8592, CST, 1:1000), p-MLKL (#37333, CST, 1:1000), t-MLKL (AP14272B, Abgent, 1:1000), GSDMD (ab209845, Abcam, 1:1000), GSDME (ab215191, Abcam, 1:1000), IRG1 (#17805, CST, 1:1000), NLRC5 (clone E1E9Y; #72379, CST, 1:2000), ASC (AG-25B-0006, AdipoGen, 1:1000), HMGB1 (Abcam, ab18256, 1:1000), LDHA (19987-1-AP, Proteintech, 1:1000). Membranes were then washed and probed with the appropriate horseradish peroxidase (HRP)-conjugated secondary antibodies (anti-mouse [315-035-047] and anti-rabbit [111-035-047], 1:5000, Jackson ImmunoResearch Laboratories). Following detection of the target protein, in some instances, membranes were stripped using Restore Western Blot Stripping Buffer (21059, Thermo Scientific), blocked with 5% skim milk, and reprobed with antibodies against additional proteins or β-actin as a loading control. Anti-β-actin antibody (sc-47778 HRP, Santa Cruz, 1:5000) was for loading control detection. Immunoblot images were acquired on an Amersham Imager using Immobilon Forte Western HRP Substrate (WBLUF0500, Millipore) or SuperSignal West Femto Maximum Sensitivity Substrate (34096, Thermo Fisher Scientific). For quantification, densitometric measurements of the target proteins were performed using ImageJ software (version 1.53k, NIH), and the measurements were normalized to the appropriate loading control.

### RNA-seq analysis of macrophages stimulated with heme plus Pam3 and heme plus TNF

WT BMDMs were stimulated in 6-well plates at a final volume of 1 mL per well with heme (50 µM) in combination with either Pam3 (500 ng/mL) for 30 h or TNF (100 ng/mL) for 32 h. Cells were homogenized in 1 mL of TRIzol Reagent (15596026, Invitrogen) and total RNA was extracted following the manufacturer’s instructions. Library preparation and next-generation sequencing were performed at the St. Jude Hartwell Center Genome Sequencing Facility using the Illumina NovaSeq 6000 sequencer. Batch 1, comprising media controls (n = 4) and heme plus Pam3-treated samples (n = 4), was sequenced using paired-end 101-cycle sequencing. Batch 2, comprising media controls (n = 2) and heme plus TNF-treated samples (n = 2), was sequenced using paired-end 151-cycle sequencing. Fastqs were aligned to mouse genome (GRCm39) using STAR^66^ with the following parameters: outFilterMultimapNmax = 1000; outFilterScoreMinOverLread = 0.25; runThreadN = 3; quantMode = GeneCounts; twopassMode = Basic.

Differential expression analysis was performed using DESeq2 v1.46.0^67^. Raw gene counts were modeled separately for each comparison using a negative binomial generalized linear model, with treatment group included in the design. Media-treated samples were used as the reference group. Genes with at least 10 counts in at least two samples were retained for analysis. Differential expression was assessed using the Wald test, and *P* values were adjusted for multiple testing using the Benjamini–Hochberg method. Genes with an adjusted *P* value < 0.05 were considered differentially expressed. Mouse MitoCarta v3.0^68^ was used to identify mitochondrial genes among the differentially expressed genes by subsetting mitochondrial genes from the genome-wide differential analysis to assess their role in heme plus Pam3 and heme plus TNF-triggered pro-inflammatory signaling. For visualization, the 25 significant MitoCarta genes with the highest log_2_ fold changes in each treatment-versus-control comparison were selected. Counts were transformed using the regularized logarithm transformation with the experimental design retained, and expression values were centered for each gene by subtracting its mean across samples. The resulting replicate-level expression values were capped at −2 and 2 for visualization and displayed without row or column clustering using the Heatmap function from the ComplexHeatmap v2.24.1^69^ in R v4.5.1. Corresponding log_2_ fold changes and adjusted *P* values were displayed alongside the expression heatmaps.

### Transcriptomic analysis of public datasets related to hemolytic conditions

Seven datasets deposited in GEO^70^ (accession IDs: GSE168532^21^, GSE241467^22^, GSE215835^23^, GSE26050^24^, GSE207625^71^, GSE83563^72^, and GSE58287^28^) were analyzed to understand the expression of *IRG1* in hemolytic conditions. These datasets represent genetic (sickle cell disease, SCD; familial hemophagocytic lymphohistiocytosis, fHLH), parasitic (malaria) and viral (dengue hemorrhagic fever, COVID-19, Ebola, and Marburg) infections. GSE168532 consists of transcriptomic profiles of classical monocytes (CD14⁺CD16⁻) isolated from the peripheral blood of 13 patients with SCD and 6 healthy donors^21^. GSE241467 comprises transcriptomic profiles of peripheral blood mononuclear cells (PBMCs) from 49 individuals in a malaria-endemic region. Samples were classified as uninfected (n = 14), asymptomatic (n = 13), or febrile with malaria (n = 22)^22^. For group-level comparisons, paired and unpaired samples within each condition were combined to generate mean expression profiles per group. GSE215835 consists of PBMC samples collected at the onset of fever from 11 study participants in Indonesia. Based on subsequent clinical outcomes, participants were classified as dengue fever (DF; n = 5) or dengue hemorrhagic fever (DHF; n = 6) to examine transcriptional differences linked to disease progression^23^. GSE26050 includes PBMC samples from 11 fHLH patients and 33 healthy controls^24^. GSE207625 includes PBMC samples from 12 patients with early-stage non-severe COVID-19. Based on clinical course, patients were classified as severe cases that progressed to a deteriorated stage (n = 6) or non-severe cases that remained non-deteriorated (n = 6), to assess differential gene expression associated with disease progression^71^. GSE83563 consists of whole-blood mRNA profiles from 22 patients infected with Ebola virus and 10 healthy adults. This dataset helped analyze transcriptomic changes associated with the hyperactive immune response characteristic of Ebola virus disease^72^. GSE58287 consists of mRNA isolated from a total of 30 PBMC samples from 15 cynomolgus macaques infected with Marburg virus. Samples were collected pre-infection (day 0, n = 15) and on various post-infection days (n = 3 per day) to track the progression of infection. This dataset allows analysis of transcriptional changes associated with Marburg virus pathogenesis, including innate immune responses over time^28^.

For each of these public datasets, quality control steps were performed, including quantile normalization using the normalize.quantiles function from the preprocessCore v1.70.0 package when counts were not normalized, followed by log_2_ transformation and calculation of mean gene expression in R (version 4.4.2). For GSE26050, the chips were processed and RMA-normalized using oligo package in R (version 4.4.2). Differences between groups or conditions were assessed using the non-parametric Kruskal-Wallis test or Mann-Whitney tests, with data analysis performed in Prism (GraphPad, versions 9.0 and 10.0).

### Immunofluorescence staining and confocal microscopy

BMDMs were seeded onto poly-D-lysine coated coverslips in 24-well plates at a density of 0.5 × 10^6^ cells/well. The stimulations were performed as indicated and described above. Cells were then washed with PBS once and fixed for 10 min at 4°C in ice-chilled 99.8% methanol (A412-4, Fisher). Cells were then washed with PBS three times (10 min each) and permeabilized and blocked by incubating in permeabilization and blocking buffer (PBS containing 10% normal goat serum (#5425, CST), 1% bovine serum albumin (A-421-10, GoldBio), and 0.5% Triton X-100 (T8787, Sigma)) for 1 h at room temperature. Cells were then incubated with the following primary antibodies overnight (16 h) at 4°C: mouse anti-ASC (clone 2EI-7, 04-147, Millipore, 1:100), rat anti-caspase-8 (clone 3B10, AG-20T-0138, Adipogen, 1:200), and rabbit anti-NLRC5 (clone E1E9Y, #72379, CST, 1:500) in permeabilization and blocking buffer. Cells were then washed with PBS three times (10 min each) and stained with secondary antibodies for 2-3 h at room temperature in the dark. The secondary antibodies used were goat anti-mouse IgG AlexaFluor-647 (#A21236, Thermo Fisher Scientific), donkey anti-rat IgG AlexaFluor-488 (#A21208, Thermo Fisher Scientific), and goat anti-rabbit IgG Alexa Fluor-568 (#A11036, Thermo Fisher Scientific). All secondary antibodies were diluted at 1:500 in permeabilization and blocking buffer. After secondary antibody staining, cells were then washed with PBS three times (10 min each) and then counterstained with 1 μg/mL DAPI (#40043, Biotium) for 5 min, washed once with PBS, and mounted using Pro-Long™ Diamond Antifade Mountant (#P36961, Invitrogen) on glass slides.

Raw images were captured by a Marianas spinning disk confocal imaging system (Intelligent Imaging Innovations) comprising a Zeiss Axio Observer inverted microscope equipped with a Yokogawa CSU-X1 spinning disk system, Prime 95B CMOS camera (Photometrics), 1.45 NA 100× oil objective, and 4 diode lasers. Slidebook software (version 6.0.24; 3i) was used for initial image capture and images were analyzed using Imaris software (version x64 9.5.1; Bitplane). Cells with specks showing colocalization of NLRC5, caspase-8, and ASC were counted per field and normalized to the total number of nucleated cells within the field of view.

### Real-time qPCR (RT-qPCR) analysis

Cells were lysed in lysis buffer, and total RNA was isolated with the PureLink RNA Mini Kit (12183018A, Invitrogen) according to the manufacturer’s instructions. The concentration of RNA was quantified with a Nano-Drop 2000 (Thermo Scientific). Then, 1 µg of total RNA was reverse transcribed with the High-Capacity cDNA Reverse Transcription Kit (4374966, Thermo Scientific) according to manufacturer’s instructions. The amplified cDNA was diluted to 10 ng/µl and RT-qPCR was performed using 40 ng of cDNA with Fast TaqMan 2 × Master Mix (4444557, Applied Biosystems) or SYBR Green 2 × Master Mix (4367659, Applied Biosystems) on a 7500 Fast Real-time PCR system (Applied Biosystems). TaqMan probe for m*Irg1* (Mm01224532_m1, 4453320, Thermo Scientific), m*Abat* (Mm00556951_m1, 4448892, Thermo Scientific), m*Idh2* (Mm00612429_m1, 4453320, Thermo Scientific), m*Slc25a2* (Mm03807525_s1, 4448892, Thermo Scientific), m*Arg2* (Mm00477592_m1, 4453320, Thermo Scientific), m*Prdx5* (Mm00465365_m1, 4448892, Thermo Scientific), h*IRG1* (Hs00985781_m1, 4453320, Thermo Scientific), *mActb* (Mm00607939_s1, 4331182, Thermo Scientific), and *18S* rRNA Endogenous Control (4319413E, Thermo Scientific) were used and SYBR Green primers for *mGls*: TTCGCCCTCGGAGATCCTAC and CCAAGCTAGGTAACAGACCCT; *mGls2*: CAGAGGGACAGGAGCGTATC and TTCTTTCGGAATGCCTGAGTC and *mAldh5a1*: CGGTCAAGGAGAGGAGCTTAC and GGACTAGCCCTCGCTTATCTTT; *mActb*: GGCTGTATTCCCCTCCATCG and CCAGTTGGTAACAATGCCATGT were used. Expression was normalized using the delta-delta cycle threshold method.

### LC–MS/MS analysis

#### Cells sample collection for targeted analysis of metabolites

After appropriate treatments, cells were collected by scraping followed by a brief centrifugation and then cells were washed once with pre-chilled 0.9% Saline buffer. Next, the cell pellets were flash-frozen in liquid nitrogen and stored at -80°C until metabolites extraction.

#### Organic acids analysis

Cell pellets were extracted with 200 µL of acetonitrile/water (1:1 v/v, pH ∼3), centrifuged at 20,000 x g for 15 min at 4°C, and then 50 µL of cleared supernatant was transferred into an autosampler vial. The extraction solution contained 10 nmol of the D6-succinate (DLM-831 Cambridge Isotope Laboratories) for every 200 µL as internal standard. 25 µL of 3-nitrophenylhydrazine (3-NPH, 200 mM in 50% acetonitrile)^73^ (N21804, Sigma-Aldrich) were added directly onto the cleared supernatant, followed by the addition of 25 µL of 1-ethyl-3-(3-dimethylaminopropyl) carbodiimide (EDC; 200 mM in 50% acetonitrile) (03449, Sigma-Aldrich) and mixed by pipetting. Next, 1.5 µL of pyridine (270407, Sigma-Aldrich) was added into the mixed solution, the tube was tightly capped and mixed by mixed by vortex and then incubated at 40°C for 30 min. The derivatization reaction was quenched by adding 100 µL of 0.2% formic acid (A117-50, Thermo Fisher Scientific) prepared in acetonitrile/water (1:1) and mixed by pipetting. The vial was placed into the LC autosampler and 4 µL were analyzed by liquid chromatography with tandem mass spectrometry (LC-MS/MS). An ACQUITY Premier UPLC System (Waters Corporation) was used for the LC separations, using a linear gradient conditions as follows: 0-0.5 min 15% B; 0.5-2.5 min 15 to 97%; 2.5-4 min 97% B; 4-4.25 min 97 to 15% B; 4.25-5 min 15% B. Mobile phase A was water supplemented with 0.1% formic acid, and mobile phase B was acetonitrile with 0.1% formic acid. The column used was an Accucore C30 (50 × 2.1 mm, 2.6 μm) (Thermo Fisher Scientific) operated at 40°C. The flow rate was 350 μl/min, and the injection volume used was 4 μl. All LC-MS solvents and reagents were of the highest purity available (water, acetonitrile, formic acid, hydrochloric acid) and were purchased from Thermo Fisher Scientific. A Xevo TQ Absolute XR Triple Quadrupole Mass Spectrometer (TQA) (Waters Corp) equipped with an UniSpray ion source was employed as detector. The TQA was operated in negative ion mode using the multiple reaction monitoring mass spec method (MRM). The MRM conditions were set to a minimum of 15 points per peak, with automatic dwell times. The operating conditions of the source were: Impactor Voltage 1.0 kV, Cone Voltage 28 V, Desolvation Temp 550°C, Desolvation Gas Flow 1000 L/h, Cone Gas Flow 150 L/h, Source Temp 150°C. Authentic organic acids standards were purchased from Sigma-Aldrich (St. Louis, MO, USA) and employed to stablish the MRM conditions. The monitored parent/daughter ions, fragmentation collision energy (CE) and the retention-time windows for each derivatized-organic acid are listed in **Supplementary Table S1**. The acquired MRM data were processed using the software application Skyline 26.1 (MacCoss Lab Software).

#### Glutamine and glutamate analysis

Cell pellets were extracted with 1 mL of methanol/acetonitrile/water (5:3:2, v/v/v) and the supernatant was then collected and dried by lyophilization. Dried extracts containing the hydrophilic metabolites were dissolved in 30 µL of water/acetonitrile (8:2, v/v) and 10 µL were used for the procedure to derivatize amino acids as described previously^74^ with some modifications. In brief, the samples were placed into glass autosampler vials and then 35 µL of sodium borate buffer (100 mM, pH 9.0) was added and mixed by pipetting. Next, 10 µL of the 6-aminoquinolyl-N-hydroxysuccinimidyl carbamate (AQC, 10 mM in acetonitrile)-derivatizing reagent (30877, Cayman Chemical) was added. The vial was sealed, mixed by vortexing and then incubated at 55°C for at least 15 min. The vial was cooled to room temperature and then 1 µL was analyzed by liquid chromatography with tandem mass spectrometry (LC-MS/MS). An ACQUITY Premier UPLC System (Waters Corp) was used for the LC separations, using a non-linear gradient conditions as follows: 0-0.4 min 3% B; 0.4-8 min 3 to 96% B (using Curve #8 of the inlet condition in MassLynx™); 8-12 min 96% B; 12-12.5 min 96 to 3% B; 12.5-14 min 3% B. Mobile phase A was water supplemented with 0.15% acetic acid, and mobile phase B was acetonitrile with 0.15% acetic acid. The column used was an Accucore C30 (50 × 2.1 mm, 2.6 μm) (Thermo Fisher Scientific), operated at 50°C. The flow rate was 300 μl/min, and the injection volume used was 1 μl. All LC-MS solvents and reagents were of the highest purity available (water, acetonitrile, acetic acid, boric acid, sodium hydroxide) and were purchased from Thermo Fisher Scientific. LC-MS/MS analysis of amino acids proceeded as described for organic acids, except that analyses were performed in positive ion mode and the Cone Voltage was set to 40 V. The monitored parent/daughter ions, fragmentation collision energy (CE) and the retention-time windows for each amino acid are listed in **Supplementary Table S2**.

### [^13^C_5_]-Glutamine tracing of itaconic acid

For isotopic labelling experiments, BMDMs were washed thrice with 1 × PBS and were stimulated appropriately in glutamine-depleted DMEM media (10313-021, Gibco) supplemented with 4 mM [^13^C_5_]-L-glutamine (HY-N0390S1, MedChemExpress), 10% heat-inactivated, dialyzed FBS (A3382001, Gibco), and 1% penicillin and streptomycin (15070-063, Thermo Fisher Scientific) for 16 h. Cells were collected by scrapping followed by a brief centrifugation, and then cells were washed once with pre-chilled 0.9% saline buffer. Next, the cell pellets were flash-frozen in liquid nitrogen and stored at −80°C until metabolite extraction.

Organic acids were extracted with 1 mL of 80:19:1 ACN/H_2_O/formic acid and 50 pmole Warfarin (Sigma-Aldrich, St. Louis, MO, USA) as internal standard. Cell pellet extractions were performed in an Omni Bead Ruptor Elite homogenizer (PerkinElmer) using 30 sec at the highest speed. Then, tubes were centrifuged at 20,000 × g at 4°C, the supernatants were transferred to new tubes, and the extracts were dried under a stream of nitrogen at room temperature. Dried extracts were resuspended with 30 µL of 89:10:1 H_2_O/methanol/formic acid, and 6 µL were used for targeted analysis of Itaconate and all its theoretical [^13^C]-isotope labelled isotopomers by mass spectrometry (LC/TQ-MRM). Briefly, an ACQUITY Premier UPLC System (Waters) was used for the LC separations, using a linear gradient condition as follows: 0–0.2 min 0% B; 0.2–2 min 0 to 97% B; 2–3.5 min 97% B; 3.5–3.6 min 97 to 0% B; 3.6–5 min 0% B. Mobile phase A is water supplemented with 0.1% formic acid; mobile phase B is acetonitrile supplemented with 0.1% formic acid. The column used was a XSelect HSS T3 (50 × 4.6 mm, 2.5 µm) (Waters) operated at 30°C. The flow rate was 600 µL/min, and the injection volume was 6 µL. All the LC-MS solvents and reagents were from the highest purity available (water, acetonitrile, methanol, formic acid) were purchased from Thermo Fisher Scientific. A Xevo TQ Absolute XR Triple Quadrupole Mass Spectrometer (TQA) (Waters) equipped with an UniSpray ion source was employed as detector. The TQA was operated in negative ion mode using the multiple reaction monitoring mass spec technique (MRM). The MRM conditions were set to a minimum of 15 points per peak, with automatic dwell times. The operating conditions of the source were: Impactor Voltage 1.0 kV, Cone Voltage 20 V, Desolvation Temp 500°C, Desolvation Gas Flow 1000 L/h, Cone Gas Flow 150 L/h, Source Temp 150°C. Authentic standard of itaconic acid was purchased from Sigma-Aldrich and used to establish the MRM and chromatographic method conditions. The monitored parent/daughters’ ions, fragmentation collision energy (CE), and the retention-time windows for data collection from all the analytes are listed in **Supplementary Table S3.** The acquired MRM data were processed using the software application Skyline 26.1 (MacCoss Lab Software). The data from theoretical daughters and all parents with [^13^C]-isotopomers were combined to calculate the presence of isotope-labelling compared to the unlabeled of itaconic acid ([M+0] peak areas).

### NAD^+^ measurement

NAD^+^ levels in cell lysates were measured using EnzyChrom NAD/NADH Assay Kit (E2ND-100, BioAssay Systems) following the manufacturer’s instructions. NAD^+^ content was normalized to total protein, which was quantified using Pierce™ BCA Protein Assay Kit (23227, Thermo Fischer Scientific).

### Ammonia measurement

Ammonia released into the conditioned media was quantified using the EnzyChrom Ammonia/Ammonium Assay Kit (ENH3-100, BioAssay Systems) following the manufacturer’s instructions.

### In vivo hemolysis model

LPS (1 mg/kg body weight; L2630, Sigma), and phenylhydrazine (PHZ; 125 mg/kg body weight; 114715, Sigma), were used for in vivo experiments. LPS was dissolved in sterile DPBS, aliquoted, and stored at -80°C until used. PHZ was freshly prepared by dissolving the weighed compound in sterile DPBS and adjusting the pH to 7.4 using 2 M NaOH, followed by filtration through a 0.22 µm syringe filter (SLGPR33RS, Millipore). Age-and sex-matched, 6-to 8-week-old WT C57BL/6J mice were injected intraperitoneally with either LPS alone, PHZ alone, or a combination of LPS with PHZ. Animals were sacrificed at 26 h post treatment, and spleen tissues and serum were collected for western blot analysis and measurement of HMGB1 and LDH levels, respectively.

### Serum HMGB1 measurement

Serum HMGB1 levels were measured using the Human/Mouse HMGB1 ELISA Kit (KE00343, Proteintech) according to the manufacturer’s instructions. Serum samples were diluted 5-fold with the sample diluent prior to analysis. HMGB1 concentrations were determined from a standard curve and expressed as ng/ml after accounting for the sample dilution.

### Serum LDH activity measurement

Serum LDH activity was measured using the CytoTox 96 Non-Radioactive Cytotoxicity Assay kit (G1781, Promega). Serum samples were diluted 20-fold in PBS. A 50 µl aliquot of diluted serum was mixed with 50 µl of LDH substrate solution and incubated for 5 min at room temperature. Stop solution was then added, and absorbance was measured at 490 nm. LDH activity was quantified using the LDH standard provided with the kit and expressed in U/ml after accounting for the sample dilution.

### Statistical analysis

Data analysis was performed using GraphPad Prism versions 9.0 and 10.0. Data are shown as mean ± SEM. Statistical significance was determined by unpaired t test for comparisons between two groups and one-way or two-way ANOVA with Dunnett’s or Tukey’s multiple comparisons tests for three or more groups. For public dataset analyses, Mann–Whitney test (two groups) or Kruskal–Wallis test (three or more groups) were applied. *P* value < 0.05 was considered statistically significant.

