## Supplementary figures and images for "Immunometabolic IRG1–NLRC5 Axis Controls Inflammatory Cell Death, PANoptosis"

### Supplemental Figure 1

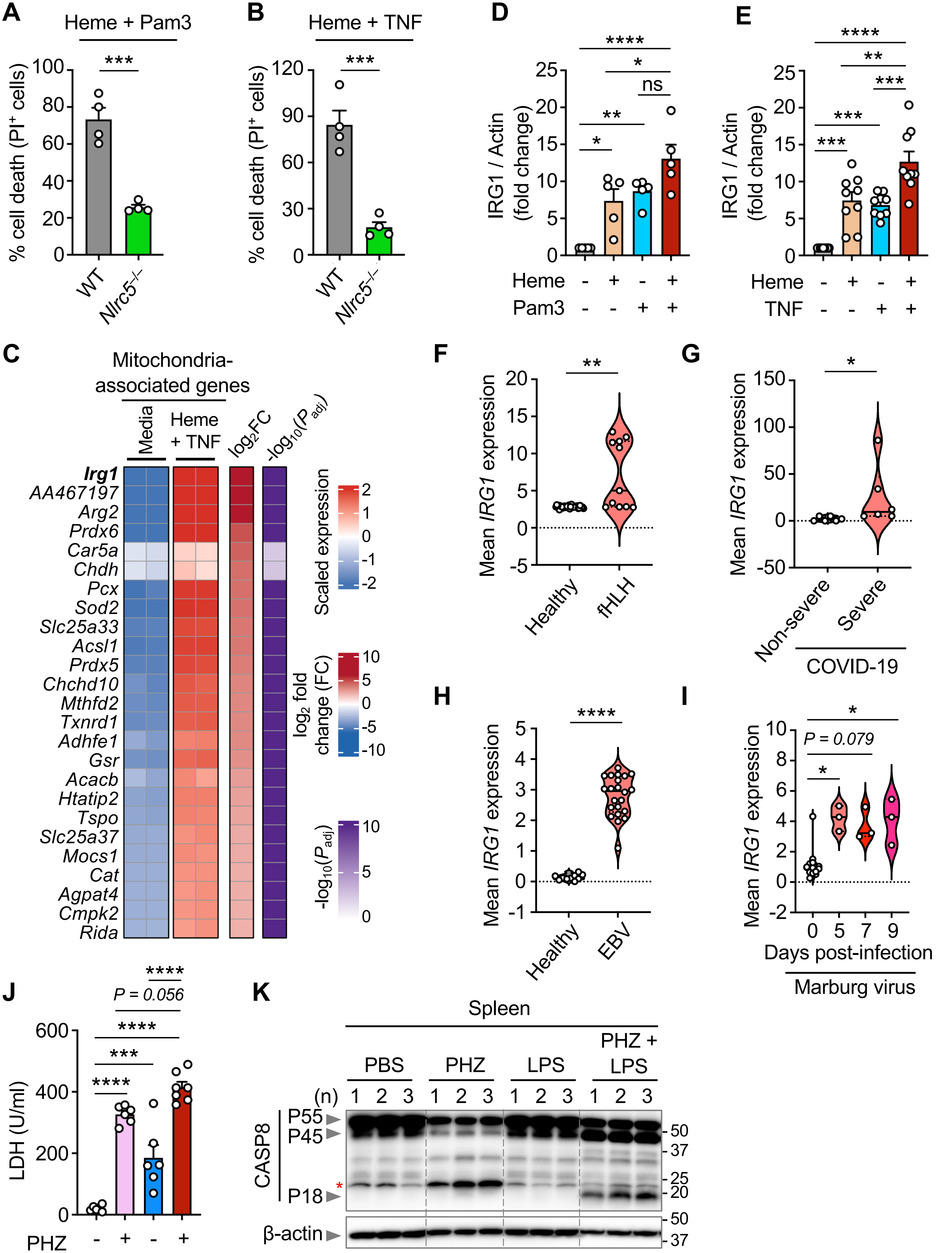


**Supplemental Figure 1**

### Supplemental Figure 2

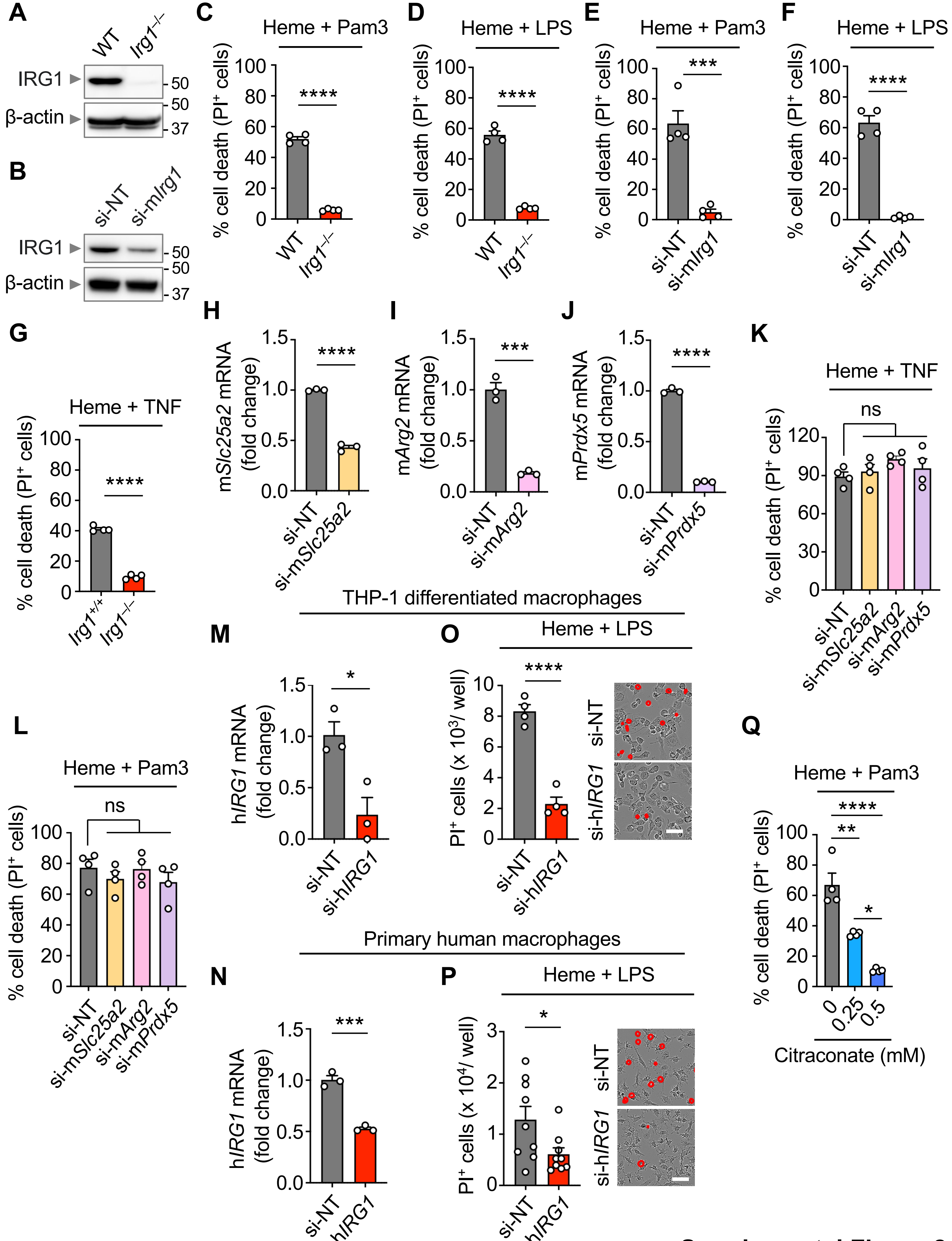


**Supplemental Figure 2**

### Supplemental Figure 3

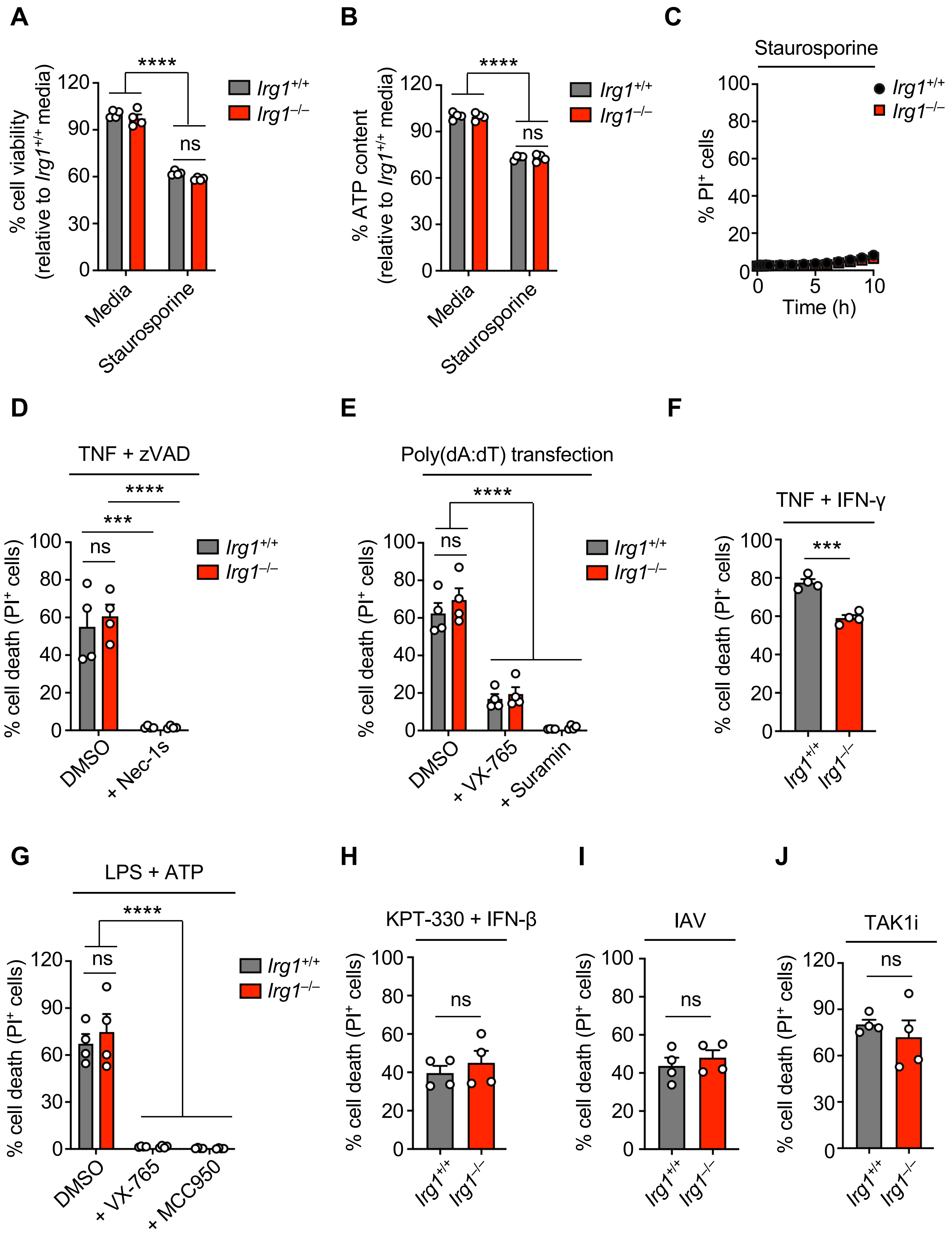


**Supplemental Figure 3**

### Supplemental Figure 4

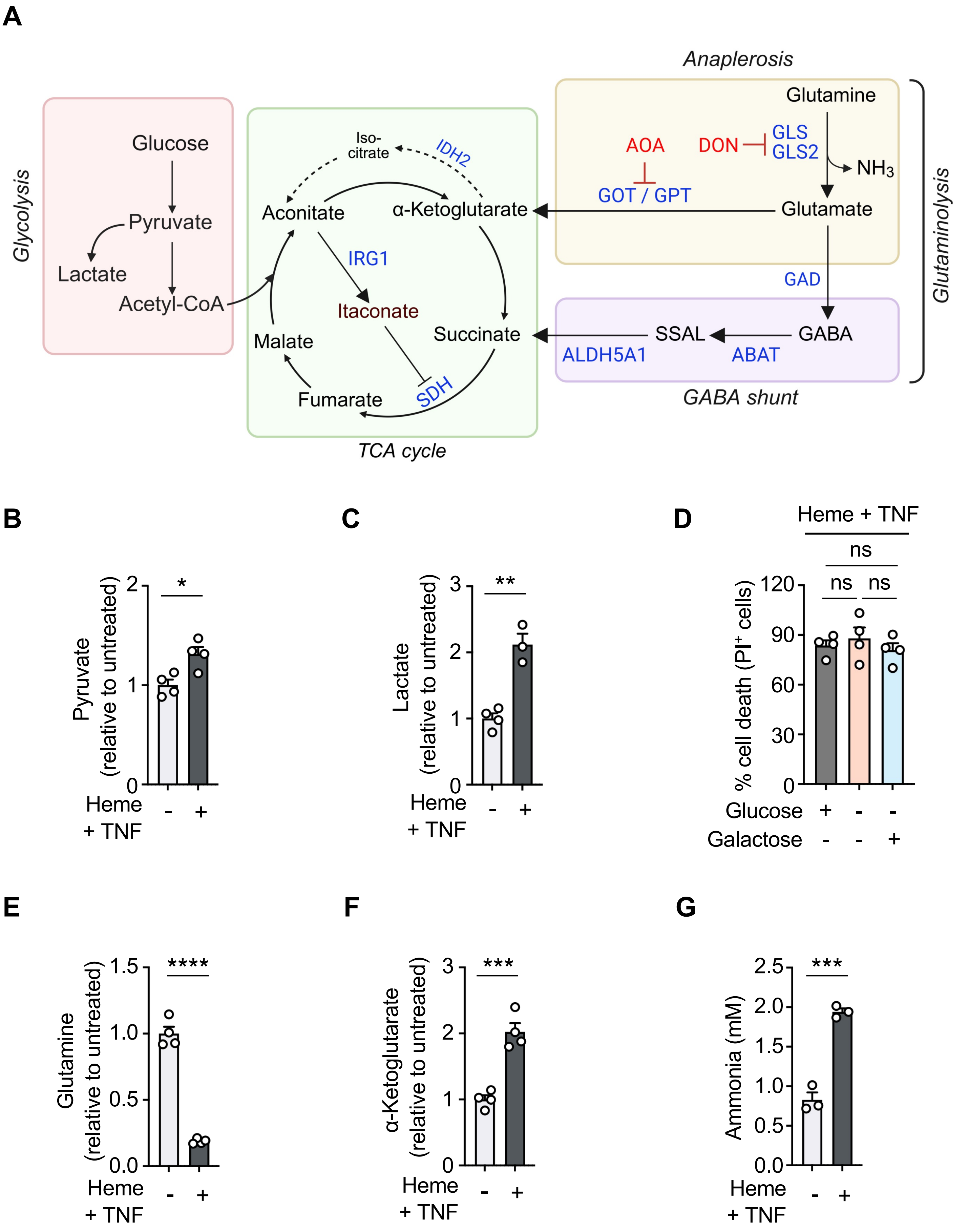


**Supplemental Figure 4**

### Supplemental Figure 5

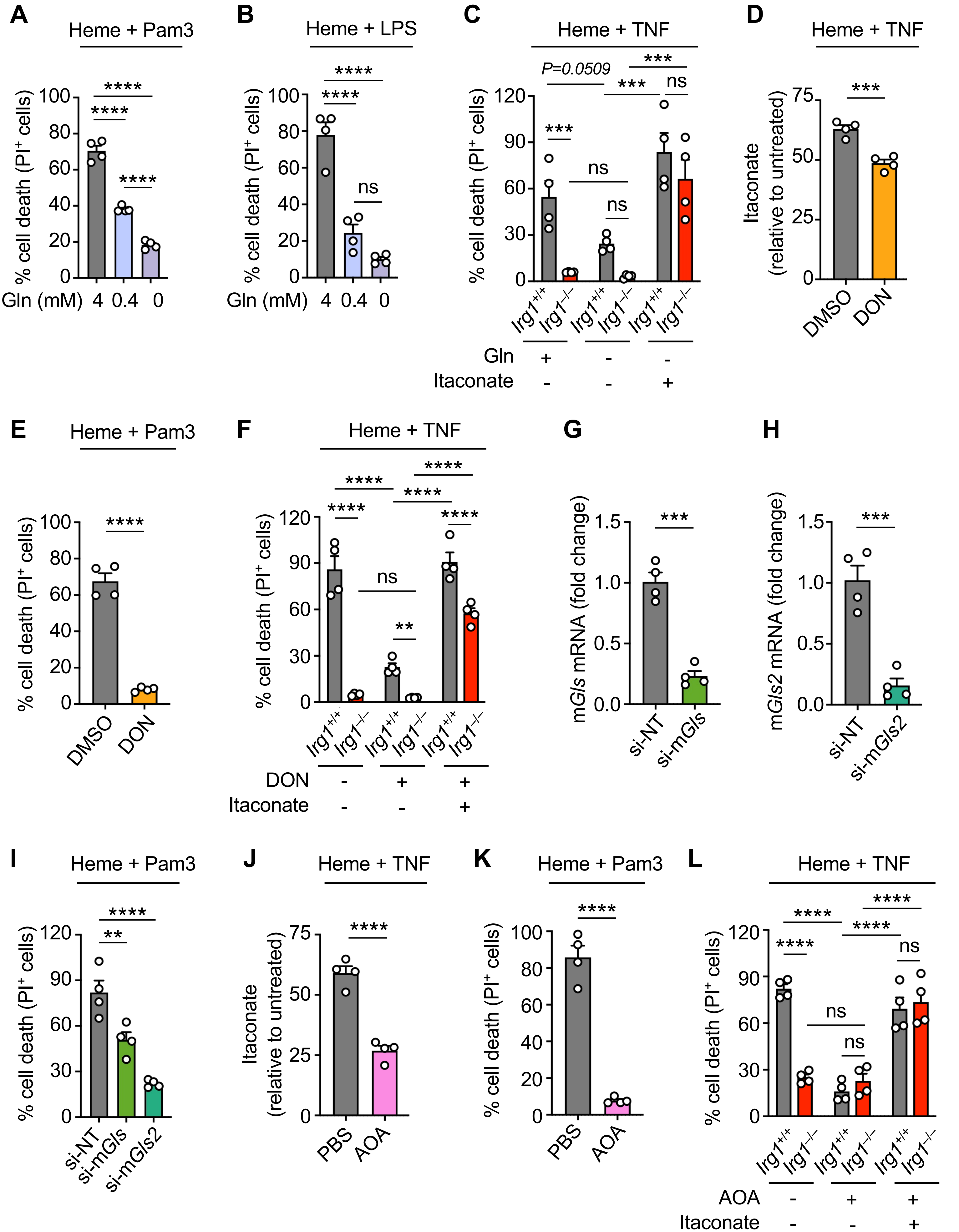


**Supplemental Figure 5**

### Supplemental Figure 6

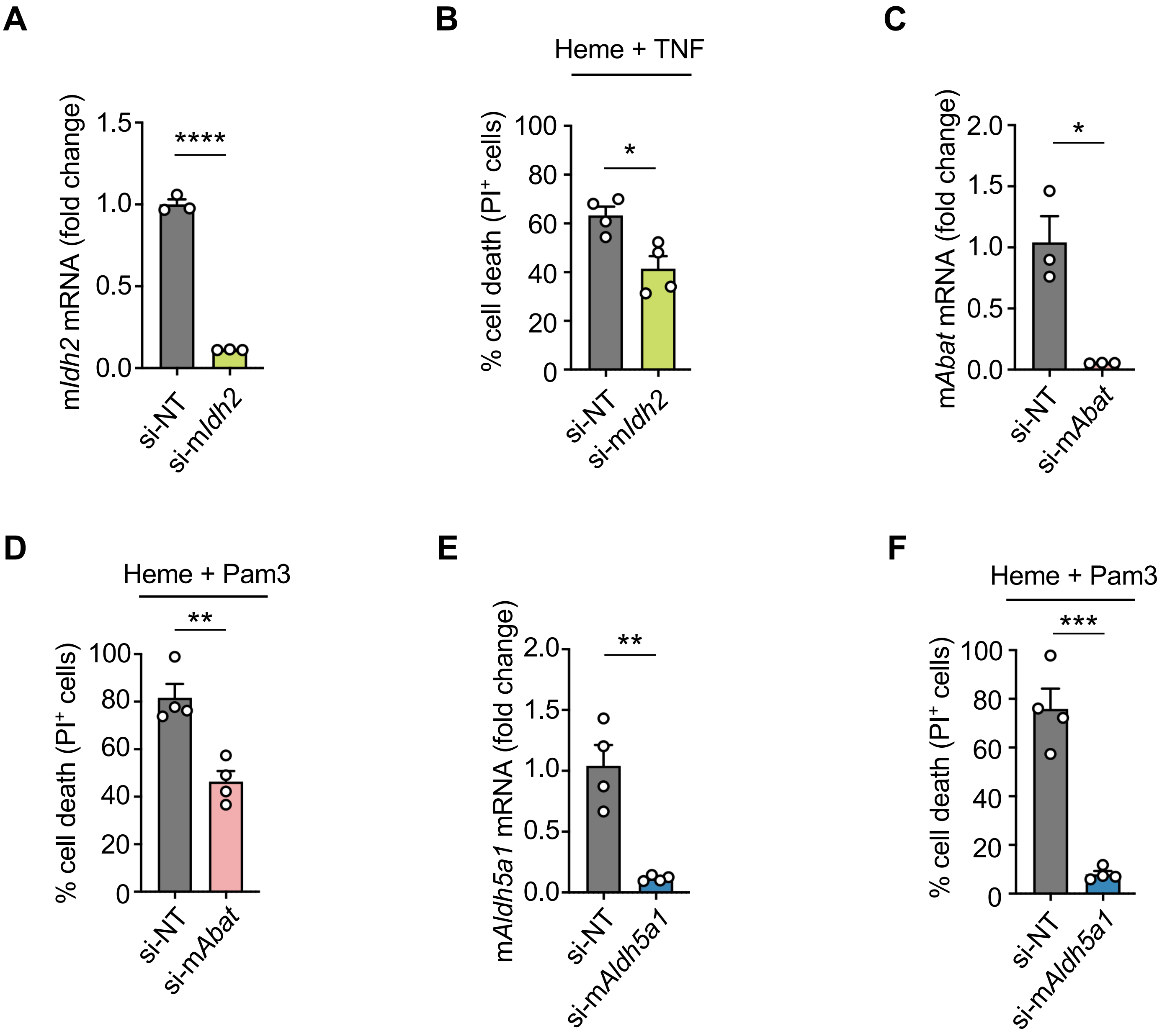


**Supplemental Figure 6**

### Supplemental Figure 7

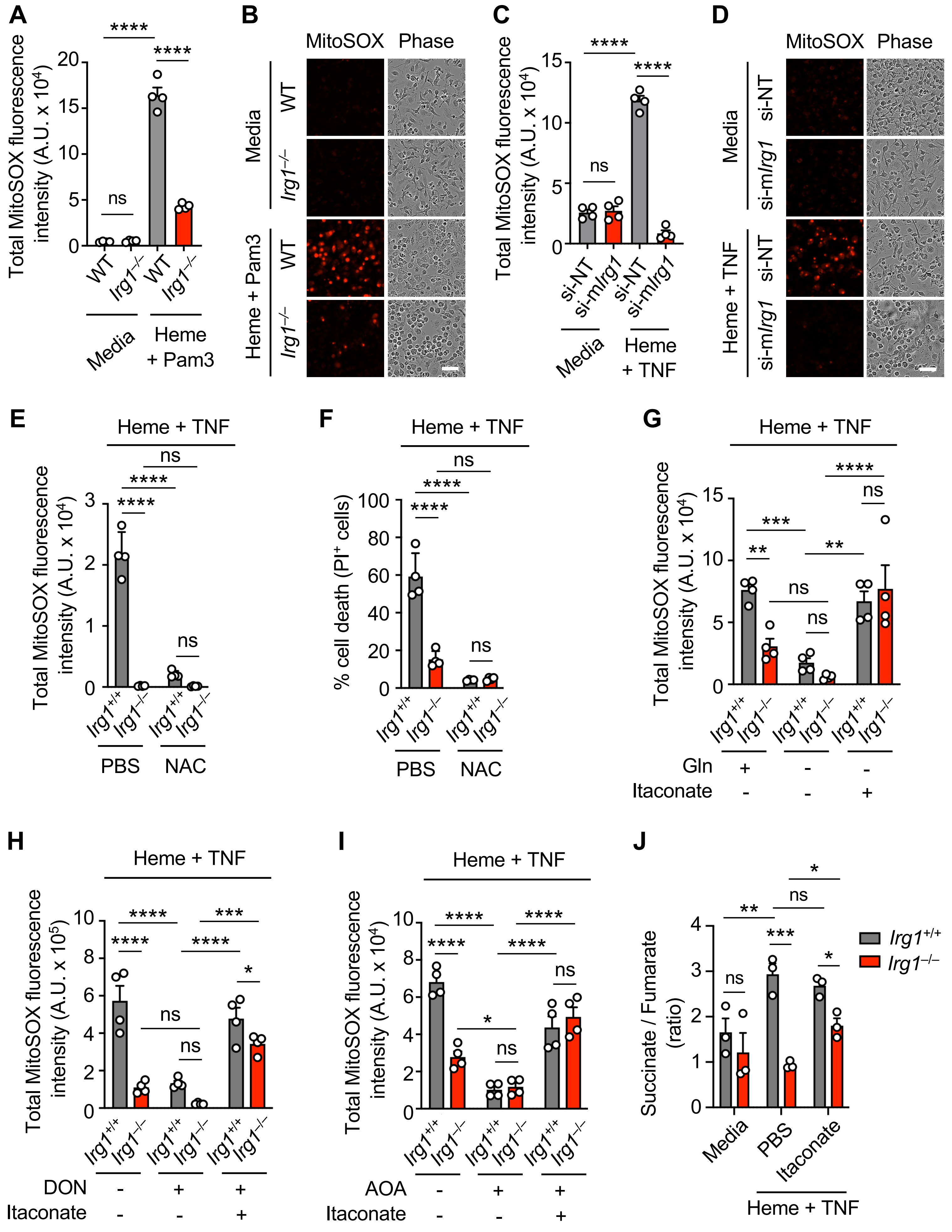


Supplemental Figure 7
