## Supplemental Table 1 for "Immunometabolic IRG1–NLRC5 Axis Controls Inflammatory Cell Death, PANoptosis"

Supplemental Table S1:

| Parent [m/z] | Daughter [m/z] | Polarity | Start [min] | End [min] | CE (eV) | Analyte |
|---|---|---|---|---|---|---|
| 223.9742 | 137.2688<br>151.9785<br>**223.9742** | Negative | 0.4 | 1.7 | 14<br>14<br>1 | Lactate-3NPH |
| 356.9304 | 136.9915<br>149.9959<br>**356.9304** | Negative | 2.4 | 3.5 | 26<br>18<br>1 | Pyruvate-3NPH |
| 385.0581 | 122.0231<br>231.5906<br>**385.0581** | Negative | 2.0 | 3.0 | 16<br>16<br>1 | Fumarate-3NPH |
| 387.0781 | 97.8969<br>234.0617<br>**387.0781** | Negative | 2.0 | 3.0 | 32<br>18<br>1 | Succinate-3NPH |
| 391.6098 | 100.6442<br>236.8253<br>**391.6098** | Negative | 2.0 | 3.0 | 34<br>16<br>1 | D6-succinate-3NPH (IS) |
| 399.0957 | 234.0603<br>246.0850<br>**399.0957** | Negative | 1.8 | 3.5 | 14<br>18<br>1 | Itaconate-3NPH |
| 403.0042 | 136.9840<br>207.9904<br>**403.0042** | Negative | 1.8 | 2.8 | 34<br>18<br>1 | Malate-3NPH |
| 550.0443 | 136.9827<br>561.0300<br>**550.0443** | Negative | 2.4 | 3.6 | 38<br>22<br>1 | $\alpha$KG-3NPH |

**In bold:** indicates the quantification ion used.
