## Supplemental Table 2 for "Immunometabolic IRG1–NLRC5 Axis Controls Inflammatory Cell Death, PANoptosis"

Supplemental Table S2:

| Parent [m/z] | Daughter [m/z] | Polarity | Start [min] | End [min] | CE (eV) | Comment |
|---|---|---|---|---|---|---|
| 316.97 | 170.93 | Positive | 0.1 | 2.5 | 30 | Gln [M+1AQC+H] |
| 318.06 | 170.93 | Positive | 0.3 | 3.5 | 30 | Glu [M+1AQC+H] |
| 308.98 | 162.86/250.97 | Positive | 7.0 | 9.0 | 14/20 | Warfarin-IS (Qty/conf) |
