## Supplemental Table 3 for "Immunometabolic IRG1–NLRC5 Axis Controls Inflammatory Cell Death, PANoptosis"

Supplemental Table S3:

| Parent [m/z] | Daughter [m/z] | Polarity | Start [min] | End [min] | CE (eV) | Analyte |
|---|---|---|---|---|---|---|
| 128.9342 | 128.9342 | Negative | 1.40 | 3.20 | 1 | Parent_M+0 |
| 130.0559 | 130.0559 | Negative | 1.40 | 3.20 | 1 | Parent_M+1 |
| 130.9516 | 130.9516 | Negative | 1.40 | 3.20 | 1 | Parent_M+2 |
| 131.9603 | 131.9603 | Negative | 1.40 | 3.20 | 1 | Parent_M+3 |
| 132.9690 | 132.9690 | Negative | 1.40 | 3.20 | 1 | Parent_M+4 |
| 133.9777 | 133.9777 | Negative | 1.40 | 3.20 | 1 | Parent_M+5 |
| 128.9342 | 41.0111 | Negative | 1.40 | 3.20 | 12 | D1_P+0 |
| 130.0559 | 41.0111 | Negative | 1.40 | 3.20 | 12 | D1+0_P+1 |
| 130.0559 | 42.0198 | Negative | 1.40 | 3.20 | 12 | D1+1_P+1 |
| 130.9516 | 41.0111 | Negative | 1.40 | 3.20 | 12 | D1+0_P+2 |
| 130.9516 | 42.0198 | Negative | 1.40 | 3.20 | 12 | D1+1_P+2 |
| 131.9603 | 41.0111 | Negative | 1.40 | 3.20 | 12 | D1+0_P+3 |
| 131.9603 | 42.0198 | Negative | 1.40 | 3.20 | 12 | D1+1_P+3 |
| 132.9690 | 41.0111 | Negative | 1.40 | 3.20 | 12 | D1+0_P+4 |
| 132.9690 | 42.0198 | Negative | 1.40 | 3.20 | 12 | D1+1_P+4 |
| 133.9777 | 42.0198 | Negative | 1.40 | 3.20 | 12 | D1+1_P+5 |
| 128.9342 | 84.9383 | Negative | 1.40 | 3.20 | 8 | D2+0_P+0 |
| 130.0559 | 84.9383 | Negative | 1.40 | 3.20 | 8 | D2+0_P+1 |
| 130.0559 | 85.947 | Negative | 1.40 | 3.20 | 8 | D2+1_P+1 |
| 130.9516 | 84.9383 | Negative | 1.40 | 3.20 | 8 | D2+0_P+2 |
| 130.9516 | 85.947 | Negative | 1.40 | 3.20 | 8 | D2+1_P+2 |
| 130.9516 | 86.9557 | Negative | 1.40 | 3.20 | 8 | D2+2_P+2 |
| 131.9603 | 84.9383 | Negative | 1.40 | 3.20 | 8 | D2+0_P+3 |
| 131.9603 | 85.947 | Negative | 1.40 | 3.20 | 8 | D2+1_P+3 |
| 131.9603 | 86.9557 | Negative | 1.40 | 3.20 | 8 | D2+2_P+3 |
| 131.9603 | 87.9644 | Negative | 1.40 | 3.20 | 8 | D2+3_P+3 |
| 132.9690 | 84.9383 | Negative | 1.40 | 3.20 | 8 | D2+0_P+4 |
| 132.9690 | 85.9470 | Negative | 1.40 | 3.20 | 8 | D2+1_P+4 |
| 132.9690 | 86.9557 | Negative | 1.40 | 3.20 | 8 | D2+2_P+4 |
| 132.9690 | 87.9644 | Negative | 1.40 | 3.20 | 8 | D2+3_P+4 |
| 132.9690 | 88.9731 | Negative | 1.40 | 3.20 | 8 | D2+4_P+4 |
| 133.9777 | 85.9470 | Negative | 1.40 | 3.20 | 8 | D2+1_P+5 |
| 133.9777 | 86.9557 | Negative | 1.40 | 3.20 | 8 | D2+2_P+5 |
| 133.9777 | 87.9644 | Negative | 1.40 | 3.20 | 8 | D2+3_P+5 |
| 133.9777 | 88.9731 | Negative | 1.40 | 3.20 | 8 | D2+4_P+5 |

Itaconic Parent (P) is a 5-carbon m/z
Itaconic Daughter 1 (D1) is a 2-carbon m/z
Itaconic Daughter 2 (D2) is a 4-carbon m/z
